# A CURE for physiological characterization of bacterioplankton in liquid culture

**DOI:** 10.1101/2020.02.21.959940

**Authors:** V. Celeste Lanclos, Jordan T. Coelho, Alex J. Hyer, Mindy M. Brooks, Emily R. Savoie, Scott Kosiba, J. Cameron Thrash

**Author notes:** Correspondence: J. Cameron Thrash, University of Southern California, Department of Biological Sciences, 3616 Trousdale Pkwy AHF 209, Los Angeles, CA 90089.

## Abstract

Bacterial characterization is an important aspect of microbiology that includes experimentally determining growth rates, environmental conditions conducive to growth, and the types of energy sources microorganisms can use. Researchers use this information to help understand and predict an organism’s ecological distribution and environmental functions. Microbiology students generally conduct bacterial characterization experiments in their coursework; however they are frequently restricted to model organisms without ecological relevance and for which the results have been known for decades. We present a course-based undergraduate research experience (CURE) curriculum to involve students in characterization of previously untested, ecologically relevant bacterioplankton cultures to identify the carbon substrates used for growth, as well as the temperature and salinity ranges conducive to growth. Students use these results to connect their organism’s physiology to the isolation environment. This curriculum also exposes students to advanced microbiology methods such as flow cytometry for measuring cell concentrations, teaches them to use the programming language R for data plotting, and emphasizes scientific communication through writing, speaking, poster creation/presentation, and social media. This CURE is an attractive introduction to scientific research and was successfully tested with 147 students during the fall semester of 2018.

## INTRODUCTION

Aquatic systems host robust bacterial communities, averaging cell densities of 10^6^ cells · ml^-1^ (1). Due to their immense population sizes, bacteria strongly influence their surrounding environments, making them important elements in ecological studies. The isolation of bacteria from their natural environment into pure culture allows for physiological characterization experiments (an important feature of microbiology research) that link physiology to ecology by elucidating growth characteristics and metabolic capabilities (2, 3). Bringing microbial characterization research into a classroom setting offers students tangible laboratory experience and the chance for deeper learning through increased engagement (4, 5, 6). Indeed, physiological characterization of bacterial isolate cultures occurs in other published curricula (7, 8) that frequently use common model laboratory isolates such as *Escherichia coli, Pseudomonas fluorescens*, or *Beneckea natriegens* (9, 10, 11). These organisms are simple to work with due to their short growth cycles, high cell density, and wealth of data with which to compare results. However, their use in a classroom laboratory setting does not provide students with the opportunity to generate new results or interact with microorganisms that have different environmental relevance. Here we present a microbiology-focused course-based undergraduate research experience (mCURE) curriculum that provides students with a realistic research experience through microbial characterization experiments in which they generate new results while also developing their written and oral scientific communication skills. Adoption of CUREs like this one into introductory biology laboratory courses can expose a greater number of students to authentic research (12) and increase the diversity of undergraduate researchers while improving STEM educational outcomes for underrepresented minorities (13).

In this curriculum, students characterize single carbon substrates used for growth, as well as the growth ranges and optima for salinity and temperature, of bacterioplankton isolates that are previously uncharacterized. The novelty of this curriculum is that, in addition to students generating new results for undescribed bacterioplankton in liquid culture, students are exposed to some of the most up-to-date techniques in the field, such as flow cytometry to track bacterial growth and data analysis using the programming language R (and an Integrated Development Environment (IDE)-RStudio) (14, 15). Students also collect and process seawater samples to gain hands-on field experience and a greater understanding of the scientific process. This mCURE curriculum can be adapted to match the flexible framework indicated in Bakshi et al. 2016 (4) and can also be utilized in series with other published courses on high throughput dilution-based isolation of bacteria (16) or the genomic characterization of bacterial isolates (17). This mCURE course has been field-tested with 147 students with the following marine bacterioplankton isolates: *Gammaproteobacteria* bacterium sp. strain LSUCC0112 (18), *Methylophilales* sp. strain LSUCC0135 (19), and a betaproteobacterial isolate in the BAL58 clade, LSUCC0117 (20). Throughout data collection and analysis, students connected physiology to ecology by contextualizing their findings to the natural environment of these isolates.

### Intended audience

The mCURE curriculum teaches students the necessary skills to use modern cultivation techniques for the characterization of BSL1 aquatic bacterial isolates in liquid culture. This includes hands-on experience with experimental design, data analysis via the programming language R, and connecting lab-generated data to a broader ecological context. The curriculum also includes exercises to improve scientific communication to expert and lay audiences and was registered as a “Communication Intensive” course with a writing focus. The intended audience for this course is first- or second-year college students majoring in a STEM field.

### Learning time

The curriculum is divided into 7 parts and can be completed in 12 weeks with one 3-hour lab period per week, but this may vary depending on the growth rates of the organisms being characterized. The isolates in the initial deployment took 7-10 days to complete their growth cycle. We recommend this project be completed in a ~15-week semester format to allow for break weeks and flexibility in scheduling. Table 1 contains the curriculum schedule without breaks.

**Table 1:**
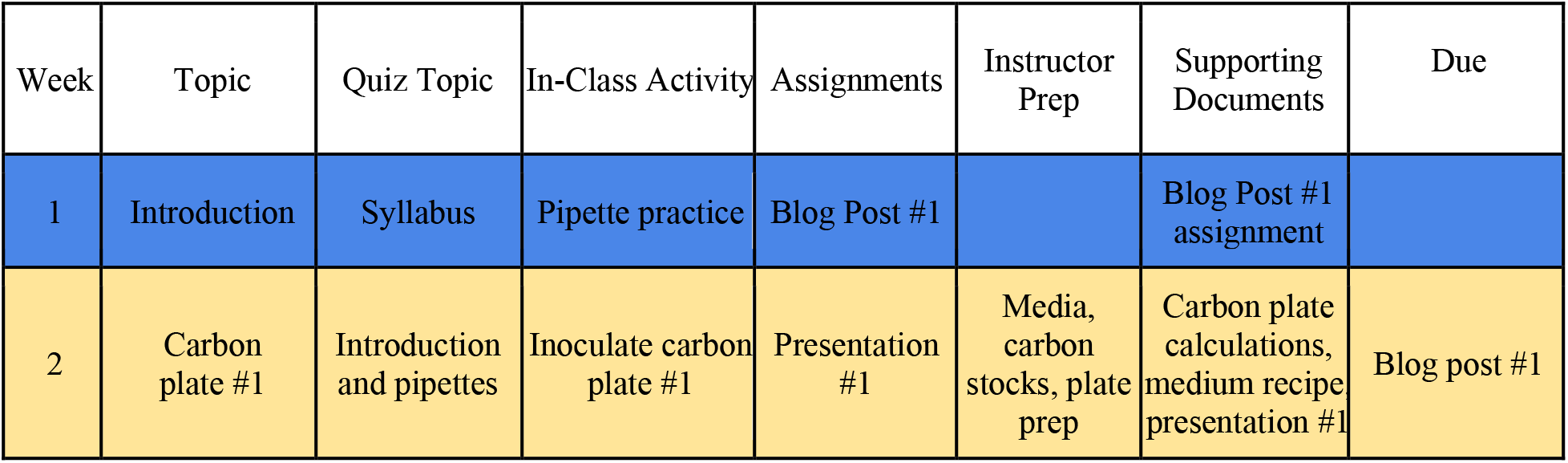
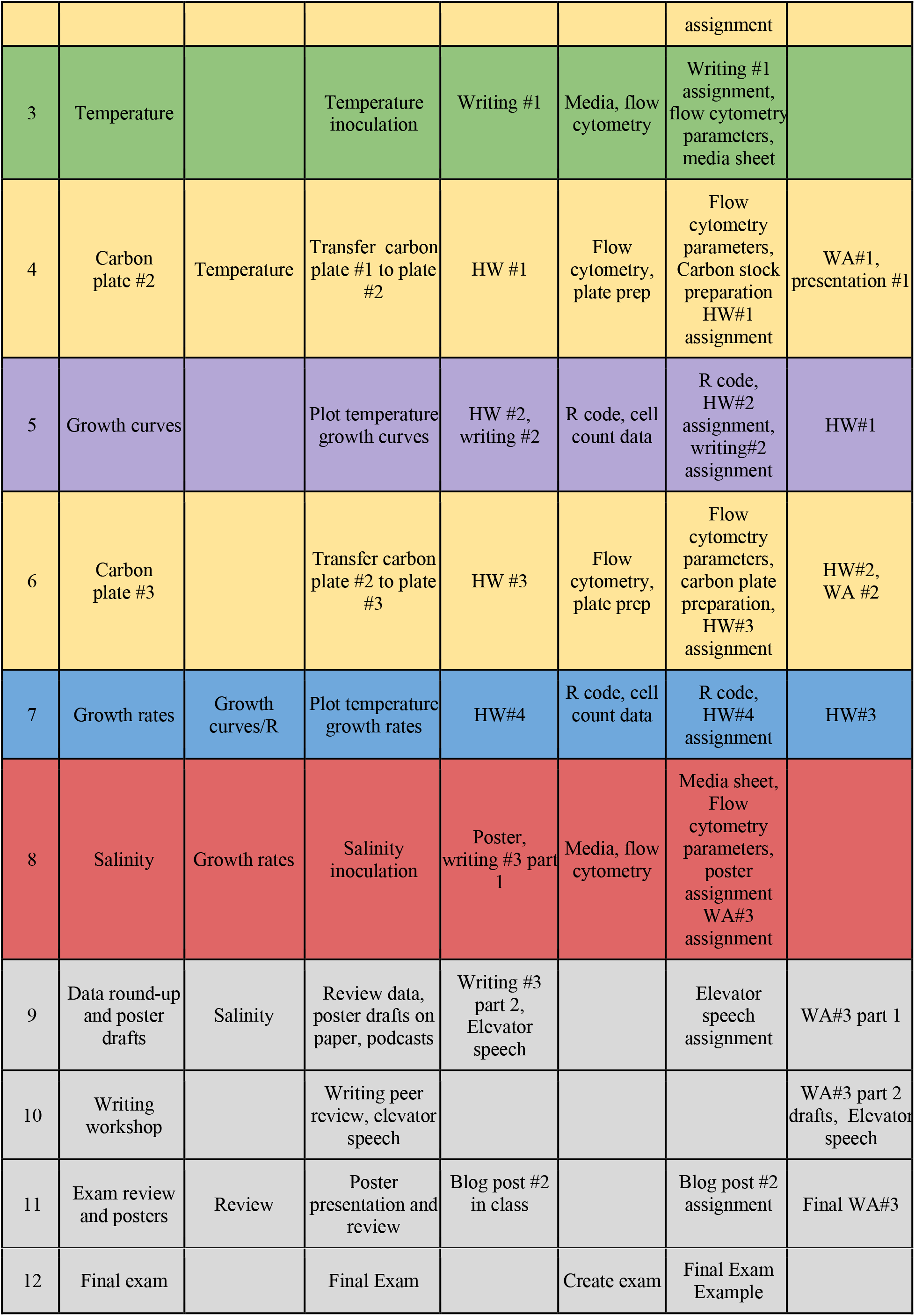
Course schedule without break weeks.

### Prerequisite student knowledge

All necessary training for students is included in the curriculum, so no prerequisites are required. However, we recommend that students have taken high school biology and chemistry.

### Learning outcomes

Upon completion of this course, students will be able to:

1. Find, read, and interpret relevant primary scientific literature;
2. Use sterile technique for proper handling of bacterial isolates in liquid media;
3. Determine physiological traits of aquatic bacterioplankton;
4. Complete basic computer scripting with R to plot growth data;
5. Link research results to publicly available ecological and environmental data;
6. Communicate research methods and results to scientific and non-scientific audiences using posters, writing, and social media.

## PROCEDURE

### Materials and overview

Cryostocks or live cultures needed for this curriculum can be obtained from a number of public culture collections, such as the American Type Culture Collection (ATCC). Pertinent cultures are also available from the Louisiana State University Culture Collection (LSUCC) and University of Southern California Culture Collection (US3C) housed in the Thrash Lab at the University of Southern California. The LSUCC was the source of the strains used in the initial curriculum deployment. To obtain cultures, please contact the corresponding author (JC Thrash). The artificial seawater medium used throughout the project has been previously published (20) and the recipe is detailed in **Appendix 1**. All media creation, experimental setup, and culture handling should be done in a biosafety cabinet to avoid contamination.

The materials needed are as follows: media components from **Appendix 1**, SYBR^®^ Green I (Invitrogen, Carlsbad, CA), a flow cytometer (e.g., the Millipore Guava easyCyte, Burlington, MA), ~20 96-well counting plates (Corning Life Sciences, Tewksbury, MA), nine 2 ml deep 96-well plates (VWR, Radnor, PA), a laminar flow hood or biosafety cabinet, ~80 125ml polycarbonate flasks (Thermo Scientific, Rochester, NY), ~10 2L Pyrex media bottles (Corning Life Sciences, Tewksbury, MA), a 10% HCl bath, and 1 set of mechanical pipettes per group (P10, P20, P100, P1000) with associated filtered tips (VWR, Radnor, PA). Lastly, students will need access to a computer for the bioinformatic portions of the course.

### Student instructions

The major parts of the semester and their corresponding week(s) and goal(s) are shown in **Figure 1**. Below we detail the experimental instructions for each segment if using an isolate from the Gulf of Mexico obtained from the LSUCC, but this curriculum can be modified for other organism types.

**Figure 1.**
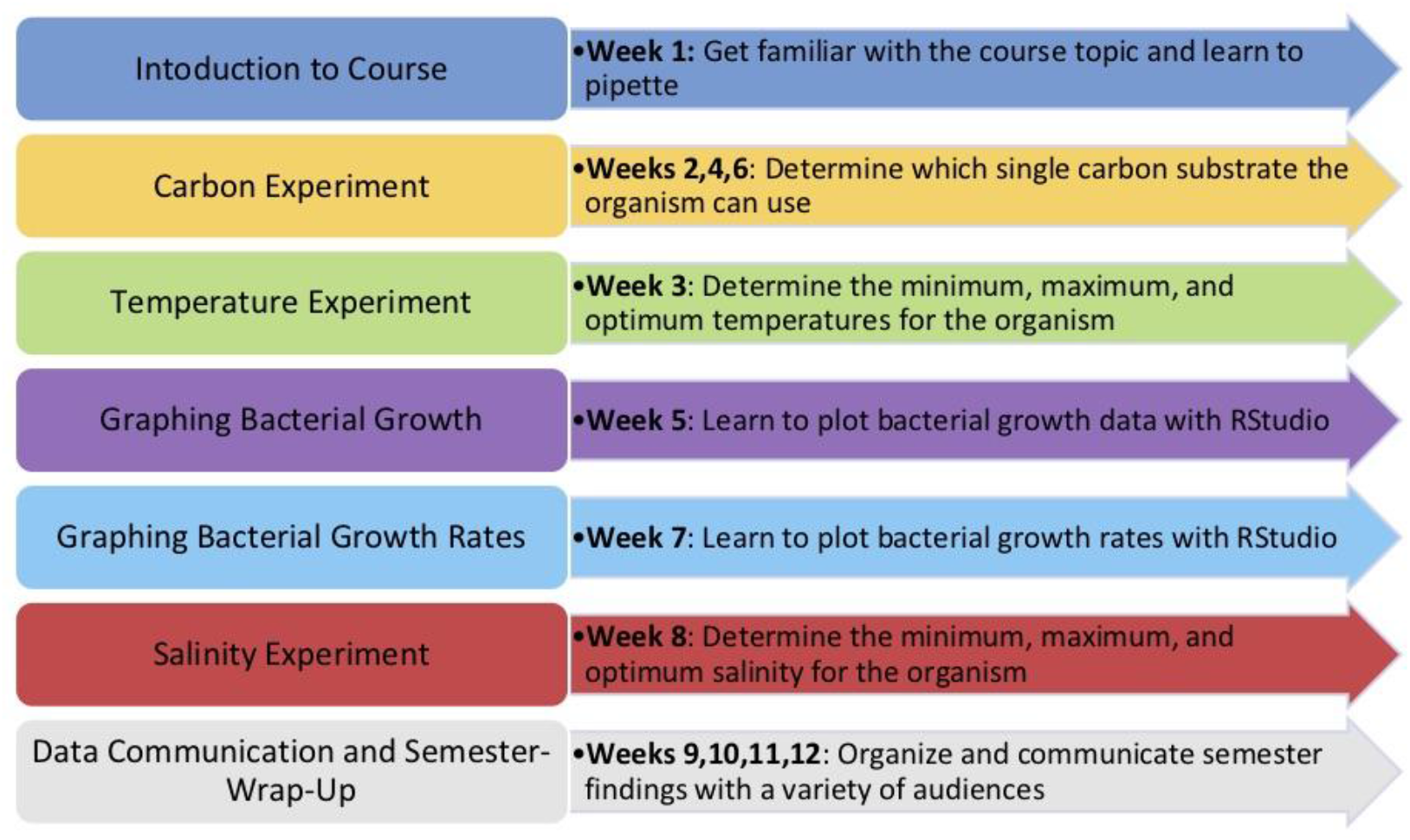
Major parts of the semester (left) with the weeks they are performed and the individual goals of the section (right).

#### 1. Introduction to the Course (Dark blue)

In the first week of the semester, students learn the structure of the course, where bacteria fit on the tree of life (21), an overview of bacterial diversity, and the instructor demonstrates how to pipette using P10, P20, and P1000 mechanical pipettes. After the first class, students write a short, informal essay highlighting their initial thoughts about their research capabilities and CURE courses (**Appendix 2**). This essay can potentially be published onto an online blog or kept for reflection at the end of the semester. Students also receive an introduction to the semester-long social media-based scientific communication assignment highlighted in **Appendix 3** and spend the last few minutes of class creating a Twitter account and sharing their handles with each other as part of this element.

#### 2. Carbon Utilization Experiment (Yellow)

This experiment uses a 96-well plate for high-throughput single carbon substrate utilization tests. The number of carbon sources per student will depend on class size with a suggested maximum of 96 total across all students to maximize the capabilities of the flow cytometer for growth quantification. This experiment has actions in weeks 2, 4, and 6 with separate instructions for incubation weeks 3 and 5 (see sections 3-4).

In week 2, students set-up carbon plate 1. Students calculate the volume of carbon stock needed to reach an appropriate concentration of carbon per well (0.5 μM for our oligotrophic organisms), as well as the volume of culture needed to inoculate a starting concentration of 1×10^4^ cells · ml^-1^ (**Appendix 4**). Using a biosafety cabinet or laminar flow hood, each student pipettes the calculated volume of their provided carbon source(s) into the designated well(s) of a 96-well plate that is pre-filled with artificial seawater medium (ASM) lacking a carbon growth substrate. Students homogenize their provided bacterial culture by swirling the culture-containing flask, then inoculate their culture into their wells that now contain ASM with a single carbon source. The plates should be covered with a sterile lid before leaving the biosafety cabinet. For organisms with a 7-10 day growth cycle, plates incubate for two weeks at isolation temperature of the strain.

In week 4, students use the flow cytometry cell count data from plate 1 provided by the instructor to calculate the volume of culture needed to inoculate a starting concentration of 1×10^4^ cells · ml^-1^ into plate 2 (**Appendix 4**). Students use a new 96-well plate pre-filled with carbonless medium and add their assigned carbon source into plate 2. Using a biosafety cabinet or laminar flow hood, students transfer the calculated volume of their cultures from carbon plate 1 to carbon plate 2, cover with a sterile lid, and let sit for the same growth period and the same temperature as plate 1. In week 6, students use the cell count data from plate 2 provided by the instructor to calculate the volume of culture needed to inoculate a starting concentration of 1×10^4^ cells · ml^-1^ into plate 3, repeating the process above for plate 2 (**Appendix 4**). After the plate 3 incubation period, students organize instructor-provided cell count data from all three plates into a results table. When cell growth is positive for a given carbon source in all three consecutive transfers, the organism is considered able to use this as a sole carbon source because the possibility that cell growth was due to carryover carbon has been eliminated.

#### 3. Temperature Experiment (Green)

In week 3, students pipette 50 mL of growth medium into their flask(s) in a biosafety cabinet or laminar flow hood, then use the cell count data from their instructor to calculate the transfer volume needed to inoculate a starting concentration of 1×10^4^ cells · ml^-1^. Flasks are then incubated at a range of ecologically relevant temperatures (e.g. organisms isolated from coastal Louisiana were placed in 12°C, 24°C, 30°C, and 40°C) for their growth cycle. Students use information from lecture about the effect of temperature on bacterial distribution and growth to predict the patterns of growth in cold versus warm temperatures that might be seen in the data and how this might relate to the ecological distribution of their organism. These predictions should be kept in their lab notebooks to compare their later results to and include in their writing assignments. Students begin their first writing assignment on bacterial carbon utilization and on creating a presentation on their assigned carbon source (**Appendices 5, 6** respectively).

#### 4. Tracking Bacterial Growth (Purple)

In week 5, students learn about bacterial growth and plot growth curves in R using data from the temperature experiments. Students are given cell count data collected by the instructor via flow cytometry (see below), download R and RStudio for homework (**Appendix 7**), and use the Code for Growth Curves (**Appendix 8**) to input their data and generate a plot of their isolate’s growth over time at various temperatures. Students connect their lab results to the data in the literature about the isolation site of the culture in their writing assignment (**Appendix 9**). Students also use other sections’ growth data (if available) to practice using R and see that organisms originating from the same general environment can have different growth preferences (**Appendix 10**).

#### 5. Calculating and Graphing Bacterial Growth Rates (Light blue)

In week 7, students evaluate poster examples and use **Appendix 11** to guide their thoughts on effective versus ineffective poster design. They also learn the components of the bacterial growth rate equation, calculate growth rate by hand (**Appendix 12**), and use Microsoft Excel to create a worksheet to calculate the experimental growth rates. They then learn how to use the growth rate R code within RStudio to display a graph of growth rates at different experimental conditions (**Appendix 13**). Students plot their rate calculations in R independently or in groups (**Appendix 14**).

#### 6. Salinity experiment (Red)

In week 8, students pipette 50 mL of their assigned salinity media into their flask(s) using a biosafety cabinet or laminar flow hood, then use the cell count data from their instructor to calculate the transfer volume needed to inoculate a starting concentration of 1×10^4^ cells · ml^-1^. Predictions about the salinities in which their cells may grow are made based on the isolation location of the bacteria and the lecture about the effect of salinity on bacterial distribution and growth. Students should attempt to connect the published salinities near the organism’s isolation site to their experiment. They should also begin thinking of the overall semester output as a single multi-part project rather than individual unrelated experiments (this is prompted in the instructions for their final writing assignment, **Appendix 15**). At this time, the final poster for the course is also assigned (**Appendix 16**). Once the data for the salinity experiment is collected, students use the R codes found in Appendices 8 and 13 to plot the growth curves and rates of their organism at multiple salinities.

#### 7. Data Communication and Semester Wrap-Up (Grey)

The final four weeks of class are focused on scientific communication and the synthesis of carbon utilization, temperature, and salinity experiments into a cohesive bacterial characterization paper. Students have an entire class period in week 9 to collaborate with each other and the instructor to ensure they generate the appropriate graphs and tables for all data from the semester. They review the semester’s data in its entirety and brainstorm poster layouts on paper. In week 10, students bring a draft of their final writing and peer edit their classmate’s work (**Appendix 15**). To practice delivering a complex biological topic to a variety of audiences, they also create an elevator speech about an outside topic of their choosing (**Appendix 17**). In week 11, students present a digital poster to their classmates for grading, have an in-class exam review, and reflect on the semester and their attitude towards research with a second informal essay (**Appendix 18**). Students take their final exam in the 12th week (**Appendix 19**). After the final exam, students should present their posters at a symposium in which members of the department are invited and encouraged to attend. See **Appendix 20** for an example of a spectator guide.

### Instructor instructions

#### 1. Introduction to the Course (Dark blue)

Prior to the start of the semester, the instructor prepares a syllabus and lecture on an overview of bacteria within the tree of life. The instructor should review lab safety rules and demonstrate proper sterile technique and proper pipetting techniques. The instructor should be trained in the dishwashing protocols (**Appendix 21**), making media (**Appendix 1**), sterile technique for bacterial cultures using a biosafety cabinet and/or laminar flow hood, and operating a flow cytometer. The flow cytometer settings we used are found in **Appendix 22**. The instructor should also familiarize themselves with how social media is used for scientific purposes to be better prepared for the social media assignment of the semester (**Appendix 3**). To better be able to predict cell count frequency and the incubation times needed, the instructor should grow the organism in its isolation medium and temperature before the start of the course.

#### 2. Carbon Utilization Experiment (Yellow)

Before the carbon experiment, the instructor picks carbon sources to test and makes the carbon stocks so they can easily be amended at environmentally relevant concentrations. The sources and concentrations for the Gulf of Mexico isolates used can be found in **Appendix 4**. Artificial seawater medium (**Appendix 1**) should be prepared without any organic carbon sources other than vitamins while excluding amino acids, miscellaneous carbon and nitrogen (C&N) mix, and fatty acids. Within a biosafety cabinet, 1.5mL of this carbonless medium should be pipetted into each well of a clean 96-well plate and provided to students in weeks 2, 4, and 6. **Appendix 4** is an example setup of carbon sources, stock concentrations, and appropriate controls for the experiment. The instructor should assign carbon sources to students, with 96 total wells across all students so that the counts can be done in a single flow cytometry plate run. For small class sizes, this means that students will have multiple carbon sources and wells that they are responsible for. In class, the instructor supervises the students’ work in the biosafety cabinet to ensure sterile technique is practiced. At the end of each plate’s incubation time, the instructor counts each well with flow cytometry and makes the data available to students for interpretation.

#### 3. Temperature Experiment (Green)

The organism’s unamended growth medium should be created before the experiment begins, with the medium and clean flasks provided to students in week 3. The instructor should assign students to temperature experiments so that there is replication in the data, such as one student with multiple flasks at a single temperature or multiple students with the same temperature. In class, the instructor supervises the students’ work in the biosafety cabinet to ensure sterile technique is being practiced. Immediately after students inoculate the flasks, the instructor obtains a to cell count, then stores the flasks at ecologically relevant temperatures such as 12°C, 24°C, 30°C, and 40°C for organisms isolated along coastal Louisiana. The instructor then obtains cell counts from the flasks at regular time points (e.g. once per day for organisms with a 7-10 day growth curve). At the end of the growth period, the instructor gathers the data into a comma-separated file (.csv) for students to plot the data in RStudio.

#### 4. Tracking Bacterial Growth (Purple)

Prior to week 4, the instructor should provide instructions on how to install the programming language R and RStudio (**Appendix 7**). They also provide students with the temperature growth data that was collected and a basic structure of the code for growth curves with annotations of the functionality of each line such as that found in **Appendix 8**. The detailed instructor version of this code is in **Appendix 23**. In class, the instructor displays RStudio with an example code and plots the measured temperature data with students. This should include a line by line explanation of what the code does and real-time troubleshooting with students as they follow along.

#### 5. Calculating and Graphing Bacterial Growth Rates (Light blue)

Prior to week 7, the instructor should gather physical or electronic examples of posters for students to evaluate in preparation of their own poster design. They also provide temperature growth rates and a basic structure of the code for students with annotations of the functionality of each line of code for growth rate plotting found in **Appendix 13**. The detailed faculty version of this code is in **Appendix 24**. In class, the instructor displays RStudio with an example code and plots the provided sample data with students. This should include a line by line explanation of what the code does and real-time troubleshooting with students as they follow along.

#### 6. Salinity experiment (Red)

Media of varying salinities that the organism might encounter in the native environment should be created before the salinity experiment begins. The instructor should assign students to salinity experiments so that there is replication in the tested conditions, such as one student with multiple flasks at a single salinity or multiple students with the same salinity. Our media is extremely adaptable-we used salinities 34.8, 23.2, 11.6, and 5.8 corresponding to medium recipes MWH 1-4 (or JW1-4) (**Appendix 1**), respectively, for isolates from the Gulf of Mexico. In class, the instructor supervises the students’ work in the biosafety cabinet to ensure sterile technique is being observed. Once the experiment has been inoculated, the instructor immediately obtains a t_0_ cell count, then again at regular time points matching that of the corresponding temperature experiment in part 4. At the end of the growth period, the instructor gathers the data into a comma-separated file for students to plot in RStudio.

#### 7. Data Communication and Semester Wrap-Up (Grey)

Prior to week 9, the instructor should gather a list of data that the students have created throughout the project. This should include all tables, growth curves, and growth rate plots. In class, the instructor helps with figure modification and provides poster feedback to the students. At the end of class in week 9, the instructor should provide an example of a scientific communication (such as a podcast or TED Talk) and guide the students in a discussion about whether the communication was effective using (**Appendix 17**).

In week 10, the instructor gives students an allotment of time to peer edit each section of a classmate’s final writing paper draft. The instructor should use the quality of the feedback given as part of each student’s grade. After peer edits in week 10, the instructor arranges students into groups in which they present their elevator pitches to each other (**Appendix 17**). If possible, students should be recorded so they can better evaluate their own communication style.

Prior to week 11, the instructor creates a grading rubric and communicates the in-class poster presentation guidelines to students (**Appendix 16**). In our experience, students should focus their presentation on the discussion and future direction sections of their posters since they generally have the same experimental data with varying interpretations and connections to larger literature.

Prior to week 12, the instructor prepares the final exam and optional practical stations (see example final in **Appendix 19**). Finally, the instructor organizes a poster symposium to highlight the students’ work. This includes reserving a space, making arrangements for printing services, setting up tables, poster boards, and providing a participation worksheet such as **Appendix 20** to guide students’ interactions as presenters and spectators.

##### Suggestions for determining student learning

Student learning can be determined through both traditional methods such as in-class quizzes (**Appendix 25**) and a final exam (**Appendix 19**), but also through communication-based assessments of learning that were highlighted and included in written formal writings (**Appendices 5, 9, 15**), peer review for formal writing (**Appendix 15**), presentations (**Appendices 6, 16-17**), and a final poster (**Appendix 16**). Assessment mechanisms in relation to the course learning objectives can be found in **Table 2**.

**Table 2:** Student learning outcomes and their respective assessments.

| Learning outcome | Assessments |
| --- | --- |
| 1. Find, read, and interpret relevant primary scientific literature | Presentation 1, Elevator speech, Final writing (**Appendices 6,17,15**) |
| 2. Use sterile technique for proper handling of bacterial isolates in liquid media | Successful completion of the protocols, Final exam practical (**Appendix 19**) |
| 3. Determine and display physiological traits of aquatic bacterioplankton | Successful completion of the protocols, Homework #2, Homework #4 (**Appendices 10 and 14**) |
| 4. Complete basic computer scripting with R to plot growth data | Successful completion of the protocols, Homework #1, Homework #2, Homework #4, Poster (**Appendices 7, 10, 14**) |
| 5. Link research results to publically available ecological and environmental data | Final writing, Poster (**Appendices 15 and 16**) |
| 6. Communicate research methods and results to scientific and non-scientific audiences using posters, writing, and social media | Poster, Elevator speech, Writing assignments, Presentations, Twitter participation (**Appendices 17,17,5,9, 15,6, 16, 17, 3**) |

##### Sample data

A carbon usage table, temperature growth curves/rates, and salinity growth curves/ rates were produced for three isolates: LSUCC0112, LSUCC0117, and LSUCC0135. LSUCC0135 and LSUCC0117 remained axenic while LSUCC0112 became contaminated sometime throughout the semester (see details below in Discussion section). Though the students had never worked with bacterial cultures prior to this course, the growth curve data exhibited good reproducibility (Figs. 2,4) as each condition included 8 replicates with one student assigned to each replicate across two sections of students. This reproducibility was seen in both isolates and in the mixed LSUCC0112 culture.

**Figure 2:**
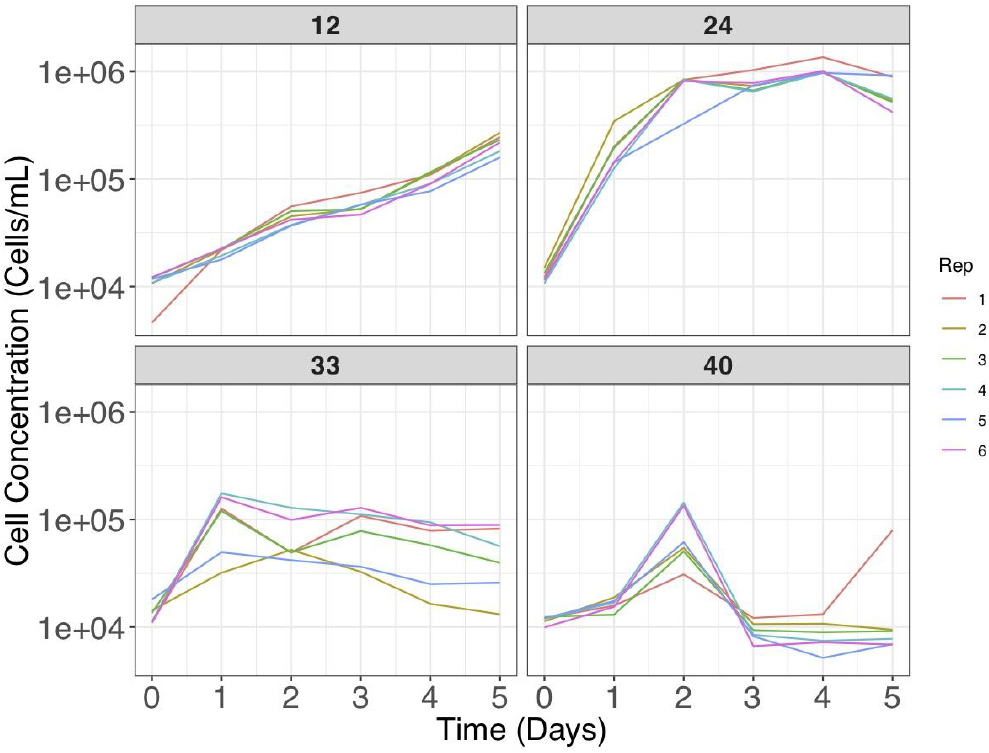
Growth curves of LSUCC0135 grown at 12°C, 24°C, 33°C, and 40°C.

Example results for each data type can be found in **Table 3** and **Figures 2–5**. LSUCC0135 had a growth temperature range of 12°C - 40°C with an optimum near 24°C. Its salinity range was 5.8 - 23.2 and the optimum salinity was undetermined but somewhere between 5.8 and 11.6. LSUCC0135 tested positive for all carbon sources after two carbon plates. LSUCC0117 had a temperature range of 12°C - 33°C with an optimum at 24°C. Its salinity range was 5.8 - 34.8 and the optimum was 11.6. LSUCC0117 could use the following carbon sources: leucine, lysine, methionine, glutamate, succinate, sucrose, serine, and folic acid (**Table 3**). The mixed LSUCC0112 culture had a temperature range of 4°C - 40°C with an optimum at 24°C. Its salinity range was 5.8 - 34.7 and the optimum was 11.5. LSUCC0112 could grow on all carbon sources at the end of carbon plate #2.

**Figure 3:**
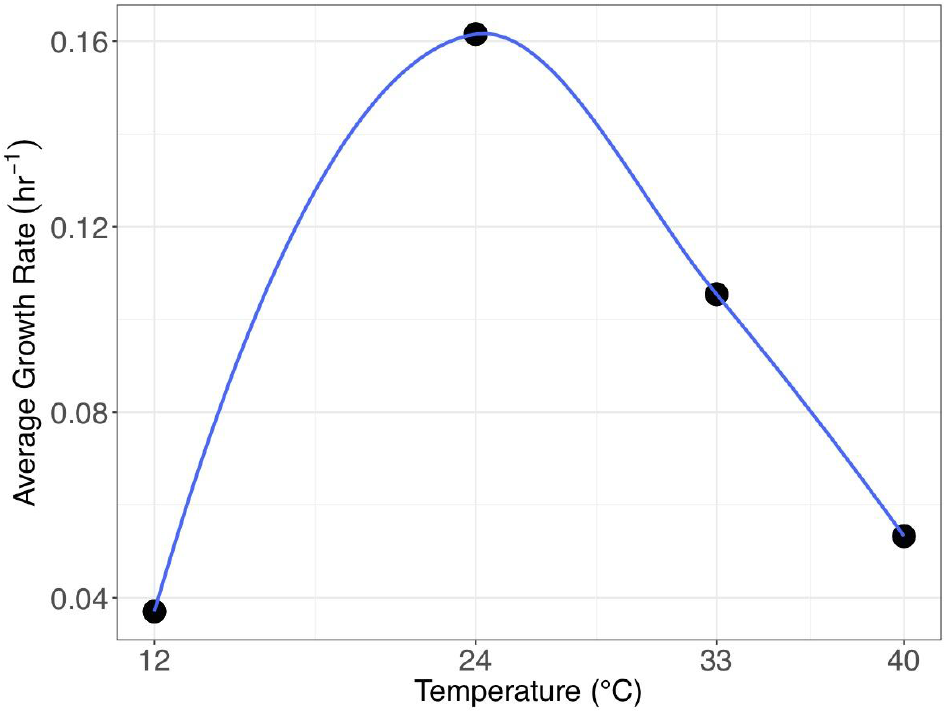
Average growth rates of LSUCC0135 grown at 12°C, 24°C, 33°C, and 40°C.

**Figure 4:**
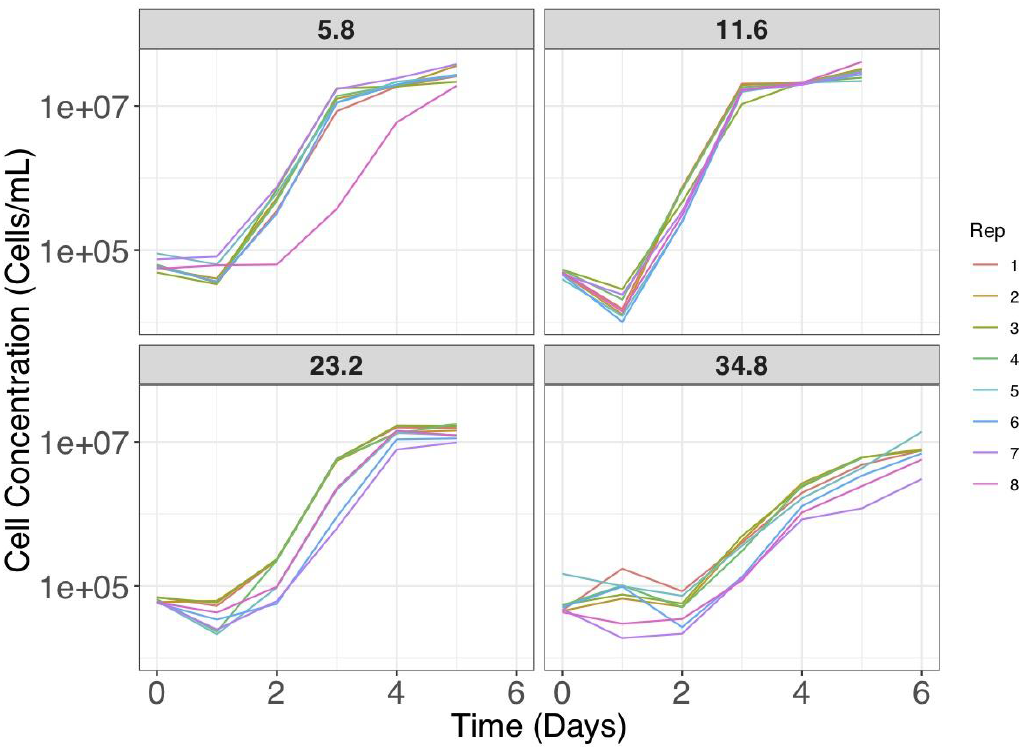
Growth curves of LSUCC0117 grown at salinities of 5.8, 11.6, 23.3, and 34.8.

**Figure 5:**
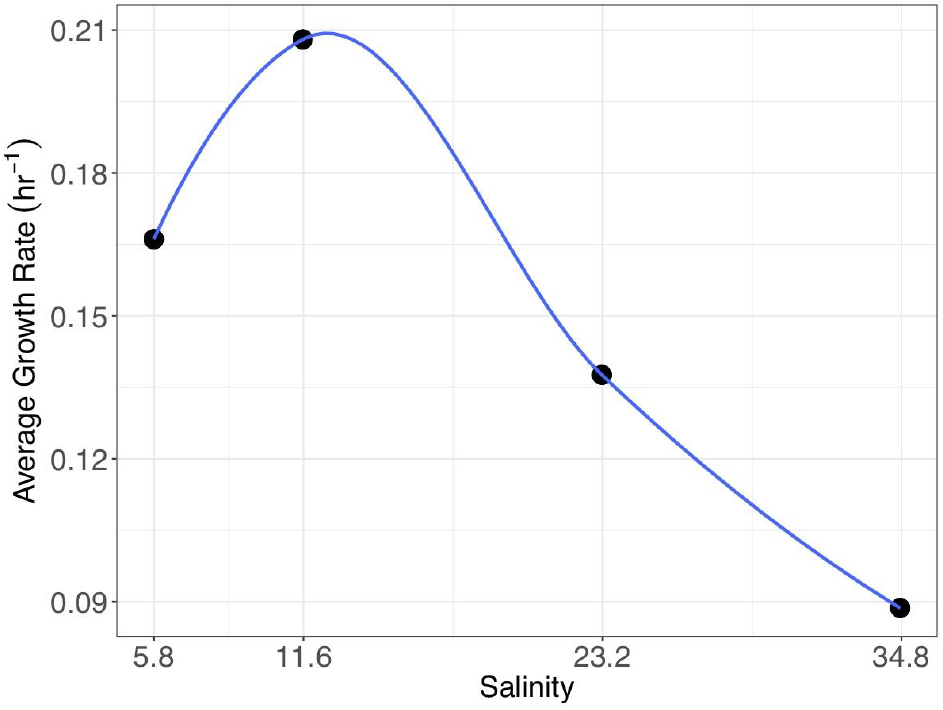
Average growth rates of 8 replicates of LSUCC0117 grown at salinities of 5.8, 11.6, 23.3, and 34.8.

**Table 3:** Example student carbon plate results for LSUCC0117.

| Carbon Source | Plate 1 | Plate 2 | Plate 3 |
| --- | --- | --- | --- |
| Threonine | NA | + | + |
| Tyrosine | NA | + | + |
| Valine | NA | + | - |
| Arginine | - | + | + |
| Cysteine | - | - | - |
| Histidine | - | - | - |
| Isoleucine | - | - | - |
| Leucine* | + | + | + |
| Lysine* | + | + | + |
| Methionine* | + | + | + |
| Phenylalanine | + | - | - |
| LSUCC0117 + ASM | + | + | NA |
| LSUCC0117 + ASM | + | + | NA |
| LSUCC0117 + ASM | + | + | NA |
| LSUCC0117 + ASM | + | + | NA |
| Glutamine | + | - | - |
| Dextrose | - | - | - |
| Ribose | - | - | - |
| Pyruvate | - | - | - |
| Citrate | - | - | - |
| Glutamate* | + | + | + |
| Acetate | - | - | - |
| Succinate* | + | + | + |
| a-ketoglutaric acid | + | + | - |
| Urea | + | - | - |
| Glycine | - | - | - |
| Choline chloride salt | - | - | - |
| Cyanate potassium salt | - | - | + |
| Sucrose * | + | + | + |
| Fructose | + | + | - |
| Glucose | + | + | - |
| Ornithine | + | + | - |
| Serine* | + | + | + |
| Folic Acid* | + | + | + |
| L-Tryptophan | - | - | - |
| LSUCC0117 + ASM-C | + | + | - |
| LSUCC0117 + ASM-C | + | + | - |
| LSUCC0117 + ASM-C* | + | + | + |
| LSUCC0117 + ASM-C | - | + | - |
| ASM-C | - | - | - |
| ASM-C | - | - | - |
| ASM-C | - | - | - |
| ASM-C | - | - | - |
| LSUCC0117 + ASM | - | + | NA |
| LSUCC0117 + ASM | - | + | NA |
| LSUCC0117 + ASM | + | + | NA |
| LSUCC0117 + ASM | + | + | NA |

\*Indicates sources that were positive for all three transfers. ASM is an abbreviation for Artificial Seawater Medium.

##### Safety issues

There are no biological safety issues in this laboratory as long as BSL1 strains are selected for investigation. If completely unknown strains are used, BSL2 protocols should be followed according to the *JMBE* Biosafety Guidelines for Handling Microorganisms in the Teaching Laboratory (22), which would also require that students be proficient in handling BSL1 strains first. Faculty should be careful with the use of glass and diluted acid for dishwashing protocols.

## DISCUSSION

### Field testing

We deployed this mCURE during the Fall 2018 semester with three graduate teaching instructors for six sections, totaling three organisms and 147 students. The two sections that characterized LSUCC0135 were in Biology 1207 Honors: Biology Laboratory for Science Majors and contained 46 students, while the other four sections working with LSUCC0112 and LSUCC0117 were in 1208 Biology Laboratory for Science Majors and contained 101 students total. Student majors included biological sciences, engineering, psychology, kinesiology, physics, animal sciences, pre-nursing, and others. Students did not know the exact project that they were signing up for during registrations, but the sections were labeled with “THIS SECTION WILL EMPHASIZE SCIENTIFIC LITERACY THROUGH PARTICIPATION IN AN UNDERGRADUATE RESEARCH PROJECT. SECTION INVOLVES COMMUNICATION-INTENSIVE LEARNING” at the time of scheduling to separate them from the more traditional curriculums.

Culture identity and purity were verified before and after the semester with 16S rRNA gene PCR (20) for the flasks that were used for experimental inoculation and we determined that LSUCC0135 and LSUCC0117 remained axenic while LSUCC0112 became mixed sometime during the semester. This could be a result of the student handling or another unknown factor.

LSUCC0135 and LSUCC0112 did not show any differential carbon usage for the first two carbon plates, and data on plate 3 was lost due to human error while an instructor transported the plates in the laboratory. Genome sequencing of LSUCC0135, however, revealed that this organism encodes RuBisCO and may be able to fix atmospheric carbon (19). This gene presence might interfere with differential carbon usage since the organism may not require organic carbon at all. The effect of this gene on heterotrophic carbon utilization would need to be further tested, but this highlights a possible complication with using uncharacterized strains for which a genome sequence is not available prior to the course.

### Evidence of student learning

Evidence of student learning is shown through the grade distribution (**Figure 6**), example poster (**Appendix 27**), example final writing excerpts (**Table 4**), and #LSUCURE on Twitter. Student grades in **Figure 2** are separated into honors and non-honors categories. Students in both categories generally did very well on posters and had a distribution of grades on the other major assignments. Students in all sections were offered up to ten bonus points throughout the semester by attending workshops about poster design or improving communication skills, but these additional points are not represented in **Figure 6**. Student presentation skills dramatically improved throughout the semester, partially due to recording their elevator speeches. Attitudes toward being on camera generally were negative at first, but students were able to watch their speech and identify nervous presentation habits. This led to more professional, confident presentations at the end of the semester with most students getting an A on their poster presentations. Videos were distributed to students, then deleted at the end of the semester to protect student privacy.

**Figure 6:**
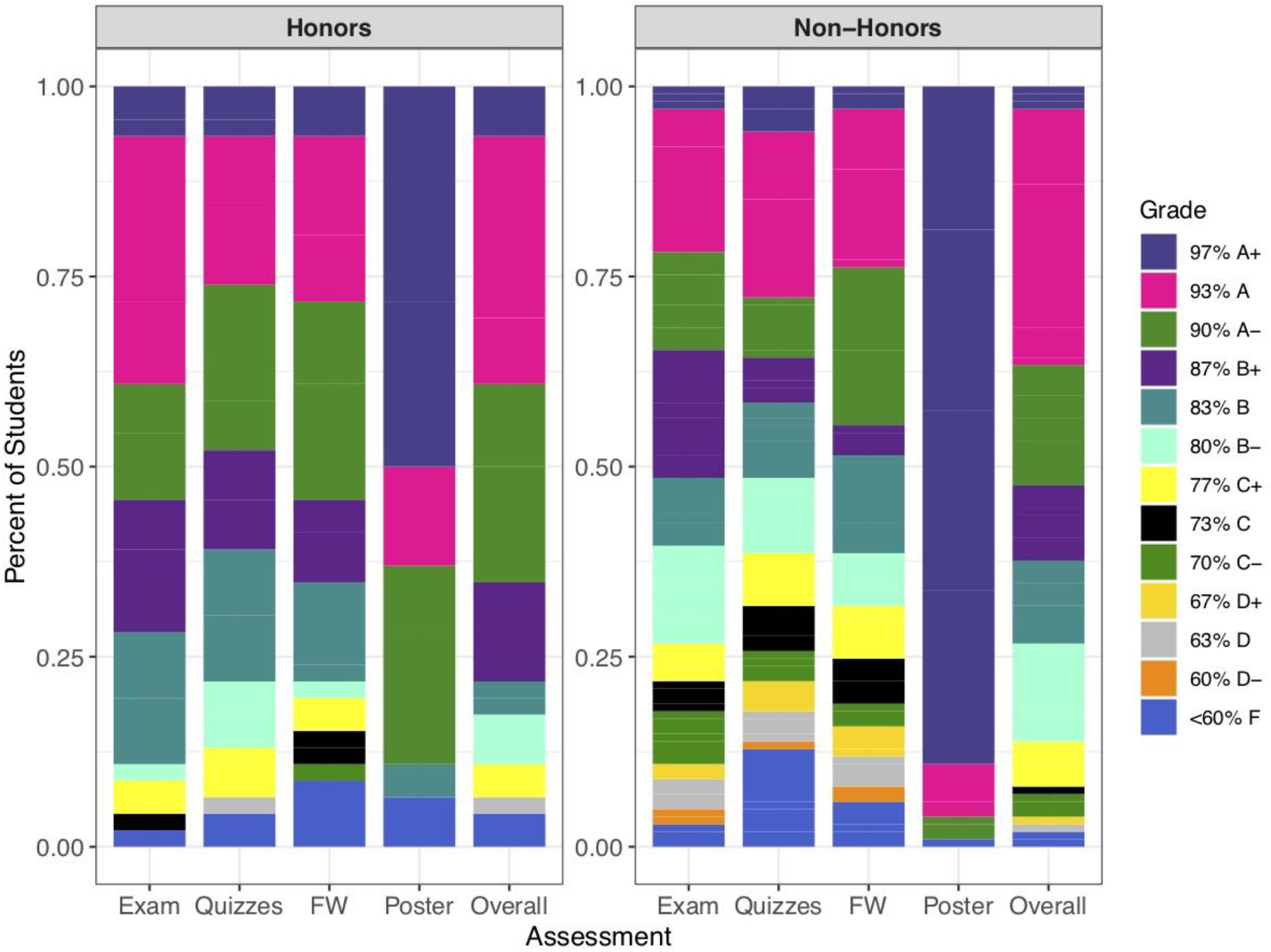
Grade distribution of the following major assignments: Final Exam (Exam), Quizzes, Final Writing (FW), Poster, and Final Grade (Overall). Data is shown for 46 students that were classified as Honors College students and 101 students that were not.

**Table 4:**
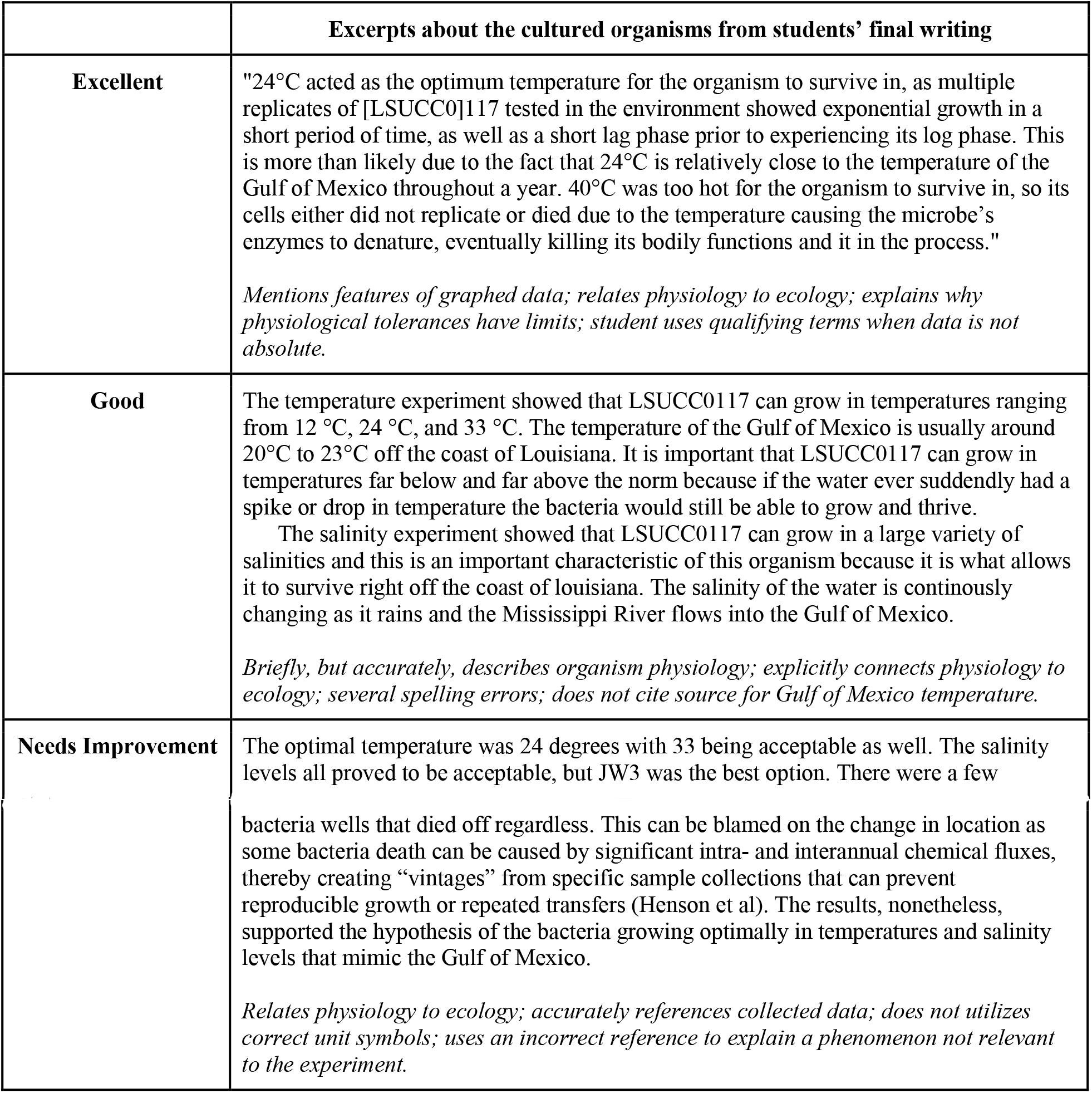
Example student writings

|  | Excerpts about the cultured organisms from students' final writing |
| --- | --- |
| **Excellent** | "24°C acted as the optimum temperature for the organism to survive in, as multiple replicates of [LSUCC0]117 tested in the environment showed exponential growth in a short period of time, as well as a short lag phase prior to experiencing its log phase. This is more than likely due to the fact that 24°C is relatively close to the temperature of the Gulf of Mexico throughout a year. 40°C was too hot for the organism to survive in, so its cells either did not replicate or died due to the temperature causing the microbe's enzymes to denature, eventually killing its bodily functions and it in the process."<br>*Mentions features of graphed data; relates physiology to ecology; explains why physiological tolerances have limits; student uses qualifying terms when data is not absolute.* |
| **Good** | The temperature experiment showed that LSUCC0117 can grow in temperatures ranging from 12 °C, 24 °C, and 33 °C. The temperature of the Gulf of Mexico is usually around 20°C to 23°C off the coast of Louisiana. It is important that LSUCC0117 can grow in temperatures far below and far above the norm because if the water ever suddenly had a spike or drop in temperature the bacteria would still be able to grow and thrive.<br>The salinity experiment showed that LSUCC0117 can grow in a large variety of salinities and this is an important characteristic of this organism because it is what allows it to survive right off the coast of Louisiana. The salinity of the water is continuously changing as it rains and the Mississippi River flows into the Gulf of Mexico.<br>*Briefly, but accurately, describes organism physiology; explicitly connects physiology to ecology; several spelling errors; does not cite source for Gulf of Mexico temperature.* |
| **Needs Improvement** | The optimal temperature was 24 degrees with 33 being acceptable as well. The salinity levels all proved to be acceptable, but JW3 was the best option. There were a few |
|  | bacteria wells that died off regardless. This can be blamed on the change in location as some bacteria death can be caused by significant intra- and interannual chemical fluxes, thereby creating “vintages” from specific sample collections that can prevent reproducible growth or repeated transfers (Henson et al). The results, nonetheless, supported the hypothesis of the bacteria growing optimally in temperatures and salinity levels that mimic the Gulf of Mexico.<br>*Relates physiology to ecology; accurately references collected data; does not utilizes correct unit symbols; uses an incorrect reference to explain a phenomenon not relevant to the experiment.* |

### Possible modifications

An attractive possible modification would be to teach students how to use the flow cytometer for growth measurements. This could only be done on a smaller class size due to the setup and run timing, the expense of the equipment, and the close supervision required for undergraduates. If the institution does not have access to a flow cytometer, another modification would be to conduct cell counts with a plate reader, via direct cell counts (microscopic counts), or via viable plate counts if the cells will grow on agar plates. This course could also be modified such that students characterize only a single organism to permit more hands-on interactions. Each section/instructor would be responsible for one portion of the characterization (carbon, temperature, or salinity) so that students can take on responsibilities of the instructor such as making media, stocks, performing cell counts, providing input on experimental design, doing dishwashing, etc. This would allow for more robust data-sharing and collaboration between students. Lastly, students could be exposed to the bioinformatic side of the workflow in a more comprehensive way by using techniques and surveys highlighted in Farrell and Carey 2018 (23).

## SUPPLEMENTAL MATERIALS

**Supplemental Table 1:**
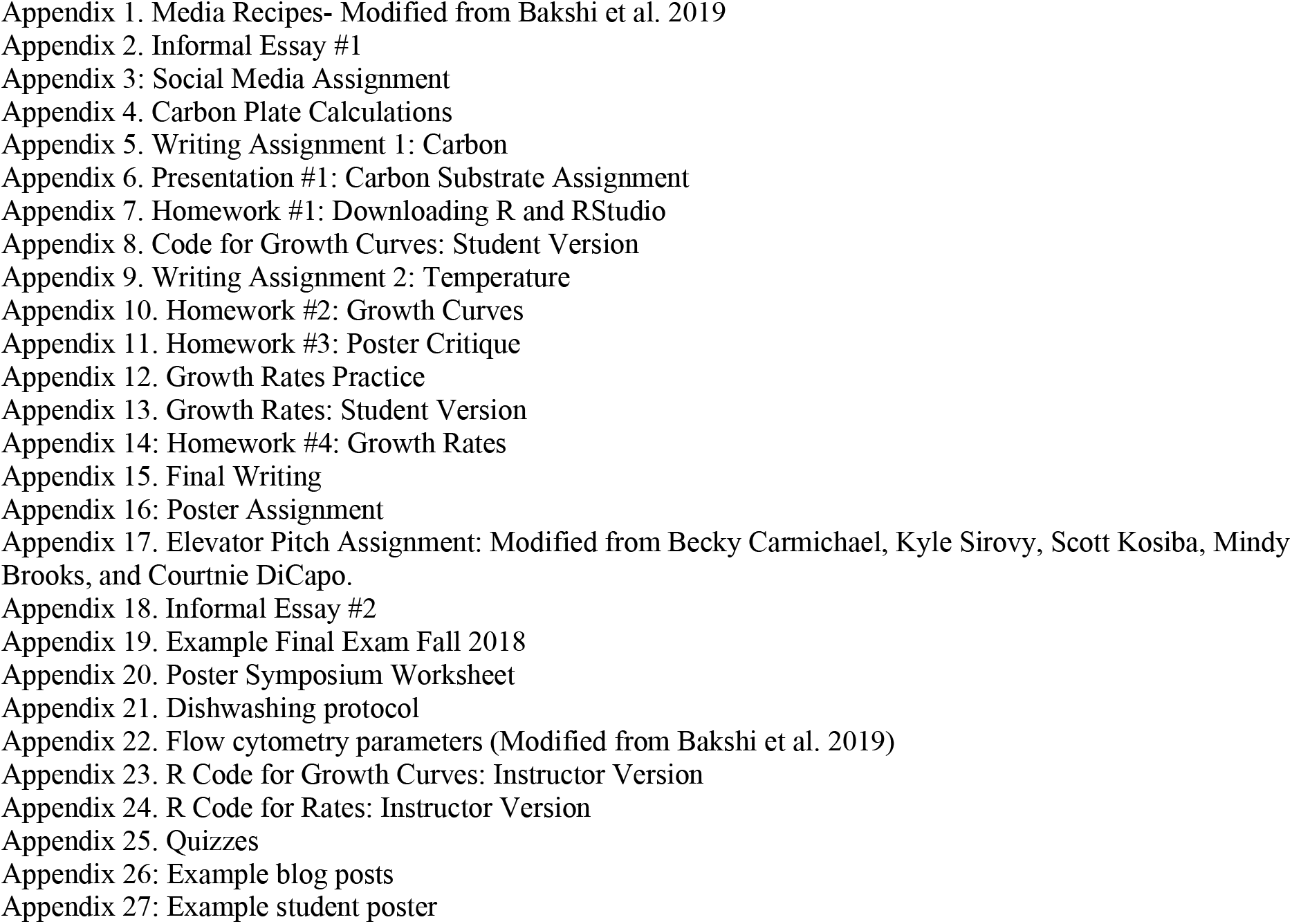
Carbon Experiment Set-Up

## ACKNOWLEDGEMENTS

We would like to thank Louisiana State University College of Science Dean Cynthia Peterson and our instructor of record, Dr. Chris Gregg, for their support of this course and the CURE program at LSU. Our undergraduate teaching assistants, Joyti Prajapati, Olivia Drago, and Zoe Long, assisted with in-class activities. Brooke Trebona helped run the poster symposium. We would also like to thank LUMCON and the crew onboard the R/V *Pelican* for facilitating an educational research cruise. Funding for this work was provided by a National Academies of Science, Engineering, and Medicine Gulf Research Program Early Career Fellowship to JCT. Lastly, thanks to the staff with LSU’s Communication Across the Curriculum (CxC) program and Becky Carmichael for her presentations about effective posters and scientific communication. Most of all, we would like to thank the LSU students that participated in this course. Their enthusiasm, drive, and feedback were instrumental in the development and success of this course.

