## Supplemental Information for "A CURE for physiological characterization of bacterioplankton in liquid culture"

**Appendix 1. Media Recipes- Modified from Bakshi et al. 2019**

Each artificial seawater medium is comprised of multiple components that are made separately and combined in the final recipe. Below we detail how to make each mix/stock, and how these are combined in the final medium. For all solutions we use acid-washed Pyrex screw-capped bottles. Note, besides the basic salts, which are made fresh for each batch of medium, the stocks and mixes can be maintained at 4˚C for additional uses. We recommend remaking the stocks and mixes every 2-3 months to avoid contamination.

**Mg/Ca stock (20x)**
In 100 mL deionized, MilliQ-filtered water, dissolve $MgCl_2$x 6 $H_2O$ (21.2 g) and $CaCl_2$x 2 $H_2O$ (3.04 g). Autoclave.

**Iron stock (1,000x)**
In 100 mL deionized, MilliQ-filtered water, dissolve $FeSO_4$x 7 $H_2O$ (0.0028 g) and Nitrilotriacetic acid (NTA) disodium salt (0.0081 g). Filter sterilize (0.2 µm).

**AA mix (50,000x)**
This can be purchased from Sigma Aldrich (Cat #M5550). Filter sterilize (0.2 µm).

**FA mix (2,000,000x)**
Combine the following: EtOH (54.86 mL), octanoic acid (15.84 mL), decanoic acid (17.26 g), isobutyric acid (9.27 mL), butyric acid (9.14 mL), and valeric acid (10.88 mL). Filter sterilize (0.2 µm).

**Inorganic N mix (2,000x)**
In 100 mL deionized, MilliQ-filtered water, dissolve sodium nitrate (0.646 g), sodium nitrite (0.028 g), and ammonium chloride (0.053 g). Filter sterilize (0.2 µm).

**Trace metals (100,000x)**
In 100 mL deionized, MilliQ-filtered water, dissolve $MnCl_2$x 4 $H_2O$ (0.018 g), $ZnSO_4$x $H_2O$ (0.002 g), $CoCl_2$(0.001 g), $Na_2MoO_4$(0.001 g), $Na_2SeO_3$(0.002 g), $NiCl_2$(0.001 g). Filter sterilize (0.2 µm).

**Vitamins (100,000x)**
In 100 mL deionized, MilliQ-filtered water, dissolve thiamine (1.69 g), riboflavin (0.003 g), niacin (0.985 g), pantothenate (1.013 g), pyridoxine (1.028 g), biotin (0.010 g), folic acid (0.018 g), B12 (0.010 g), myo-inositol (0.901 g), and 4-aminobenzoic acid (0.823 g). Filter sterilize (0.2 µm). Wrap container in foil to avoid photodegradation of vitamins.

**P mix for MWH-type media (1,000x)**
In 100 mL deionized, MilliQ-filtered water, dissolve orthophosphate (0.0022 mL) and $KH_2PO_4$(0.068 g). Filter sterilize (0.2 µm).

**Misc mix for MWH-type media (20,000x)**
In 100 mL deionized, MilliQ-filtered water, dissolve L-glutamine (0.146 g), dextrose (0.180 g), D-ribose (0.150 g), sodium pyruvate (0.110 g), sodium citrate (0.294 g), oxaloacetic acid (0.132 g), Sodium acetate (0.082 g), Sodium succinate (0.162 g), alpha-ketoglutaric acid (0.168 g), urea (0.606 g), glycerol (0.074 mL), glycine betaine (0.154 g), choline (0.140 g), sodium thiosulfate (0.158 g), cyanate (0.003 g), DMSO (0.056 mL), and DMSP (0.011 g). Filter sterilize (0.2 µm).

**Misc mix JW-type media (2,000x)**
In 100 mL deionized, MilliQ-filtered water, dissolve L-glutamine (0.015 g), dextrose (0.018 g), D-ribose (0.015 g), sodium pyruvate (0.011 g), sodium citrate (0.029 g), oxaloacetic acid (0.013 g), Sodium acetate (0.008 g), Sodium succinate (0.016 g), alpha-ketoglutaric acid (0.017 g), urea (0.061 g)

**Basic salts:**

| | JW1 | JW2 | JW3 | JW4 | MWH1 | MWH2 | MWH3 | MWH4 |
|---|---|---|---|---|---|---|---|---|
| H2O (mL) | 950 | 967 | 987.250 | 991.5 | 950 | 967 | 987.250 | 991.5 |
| NaCl (g) | 23.840 | 15.901 | 7.939 | 3.969 | 23.840 | 15.901 | 7.939 | 3.969 |
| KCl (g) | 0.746 | 0.498 | 0.248 | 0.124 | 0.746 | 0.498 | 0.248 | 0.124 |
| NaHCO3 (g) | 0.840 | 0.840 | 0.840 | 0.840 | 0.840 | 0.840 | 0.840 | 0.840 |
| Na2SO4 (g) | 4.270 | 2.848 | 1.422 | 0.711 | 4.270 | 2.848 | 1.422 | 0.711 |
| NaBr (g) | 0.082 | 0.055 | 0.027 | 0.014 | 0.082 | 0.055 | 0.027 | 0.014 |
| H3BO3 (g) | 0.026 | 0.017 | 0.009 | 0.004 | 0.026 | 0.017 | 0.009 | 0.004 |
| SrCl2 (g) | 0.014 | 0.009 | 0.005 | 0.002 | 0.014 | 0.009 | 0.004 | 0.002 |
| NaF (g) | 0.002 | 0.002 | 0.001 | 0.001 | 0.002 | 0.002 | 0.001 | 0.001 |
| KH2PO4 (g) | 0.007 | 0.007 | 0.007 | 0.004 | X | X | X | X |

**Final recipe-** In a biosafety cabinet, add the following to the basic salts mix while stirring and filter sterilize (0.1 µm) the entire medium into a sterile container. Check medium pH, which should be ~8.2-8.3. Wrap with foil. Store at room temperature prior to dispensing.

| Mix | JW1 | JW2 | JW3 | JW4 | MWH1 | MWH2 | MWH3 | MWH4 |
|---|---|---|---|---|---|---|---|---|
| Mg/Ca (mL) | 50 | 33 | 17 | 8.5 | 50 | 33 | 17 | 8.5 |
| Iron (mL) | 1 | 1 | 1 | 1 | 1 | 1 | 1 | 1 |
| AA Mix (µL) | 20 | 20 | 20 | 20 | 20 | 20 | 20 | 20 |
| FA Mix (µL) | 0.5 | 0.5 | 0.5 | 0.5 | 0.5 | 0.5 | 0.5 | 0.5 |
| Inorganic N (mL) | 0.5 | 0.5 | 0.5 | 0.5 | 0.5 | 0.5 | 0.5 | 0.5 |
| Trace Metals (µL) | 10 | 10 | 10 | 10 | 10 | 10 | 10 | 10 |
| Vitamins (µL) | 10 | 10 | 10 | 10 | 10 | 10 | 10 | 10 |
| Misc Mix JW (mL) | 0.5 | 0.5 | 0.5 | 0.5 | X | X | X | X |
| Misc Mix MWH (µL) | X | X | X | X | 50 | 50 | 50 | 50 |
| P Mix (mL) | X | X | X | X | 1 | 1 | 1 | 1 |

For additional media recipes that use modifications of the basic salts solution, see:

Henson, Michael W., V. Celeste Lanclos, Brant C. Faircloth, and J. Cameron Thrash. 2018. **Cultivation and genomics of the first freshwater SAR11 (LD12) isolate.***The ISME Journal* 12:1846-1860

**Appendix 2. Informal Essay #1**

In paragraph format, write an informal essay that highlights your initial impression about what it means to take part in a Course-based Undergraduate Research Experience (CURE) lab. There is no strict word count on this assignment, but your goal should be about half a page.

Here are some topics to cover:

-Background about yourself such as your major, goals you hope to achieve with your degree, interest (or lack of) in research, previous exposure (or lack of) to research, and conceptions about who does research

-What you've heard about CURE labs (if anything), your worries about the course, excitement towards the course, and what you hope to get out of this course

-Do you think that research experience applies to you and your goals? Why or why not?

**Appendix 3: Social Media Assignment**

Purpose: The Social Media Assignment is designed to get you engaged with the scientific community on Twitter — which many biologists have chosen as their preferred form of rapid scientific communication. By following scientists on Twitter, you will be exposed to up-to-date research articles, current interests of the scientific community, and ongoing scientific events such as conferences. Many biologists tweet articles they've recently read or talks they're currently attending and find interesting. Finally, you will need to perform some outreach of your own.

Week 1: Sign up for a Twitter account at https://twitter.com and follow your TA. Each TAs Twitter handle can be found in the syllabus. Complete the assignment on Moodle for Week 5 by telling your TA your Twitter handle so they know who to actually accept and/or follow back.

- Feel free to make your Twitter account private, but you will have to follow a few people throughout the semester.
- Your TA will not necessarily follow you back, you following them is a confirmation step. Class- relevant communication will take place over Moodle, Email, or potentially Slack, but not Twitter.
- One exception: if your Twitter account is marked as "protected", your TA will have to follow you to add you to their list. You must add them back to receive credit.

Week 2: Follow a scientist at your university and one from a different university on Twitter. Retweet a science-related tweet from each that you like, with a comment about why you liked the tweet (you cannot simply retweet, it *must* be a "Retweet with comment"). Submit the following for each scientist:

1. The name of each scientist you are following and the university or institution where they work.
2. A screenshot of your "Following" list showing that you are, in fact, following said scientists.
3. A link to, or screenshot of, your retweet (link preferred, links may not work if your account is protected).

Week 3: Follow a scientific organization on Twitter. Visit their website and tweet a sentence or two about the organization and include their web page. Submit the following:

1. The name of the organization you're following.
2. A screenshot of your "Following" list showing that you are, in fact, following said organization.
3. The link to the web page of the organization
4. A link to, or screenshot of, your retweet (link preferred).

Week 4: Follow a scientific government department, agency, etc. on Twitter. Retweet a science-related tweet from them and comment on the tweet. Suggestions include NASA, the USGS, DoE, etc. Submit the following:

1. The name of the agency you're following.
2. A screenshot of your "Following" list showing that you are, in fact, following said agency.
3. A link to, or screenshot of, your retweet (link preferred).

Week 5: Tweet about something you learned in any of your science classes this week that you found interesting or surprising. Be sure to include the course name in your tweet. Submit the following:

1. The name of the course.
2. A link to, or screenshot of, your tweet (link preferred).

Week 6: Tweet a sentence or two about a scientific news story (*not* a research article). Please avoid clickbait stories and stick to articles with primary sources as references. Submit the following:

1. The title of the story.
2. A link to the story.
3. A link to, or screenshot of, your tweet (link preferred).

Week 7: Tweet a sentence or two about a scientific video you watched and include the link to the video. It can be from any source such as YouTube, a news site, a university website, etc. As with last week, avoid clickbait and stick to videos about real science. This video needs to have taught you something new or easily described a concept you've previously found difficult. Submit the following:

1. The title of the video.
2. A link to the video.
3. A link to, or screenshot of, your tweet (link preferred).

Week 8: Tweet a sentence or two about your lab work thus far. Focus on skills you've learned or anything interesting that is happened. Submit the following:

1. A link to, or screenshot of, your tweet (link preferred).

Week 9: Choose a scientific paper tweeted byany scientist you're following and read the abstract (you do *not* have to read the whole article). Retweet the article and write one sentence about the abstract (you don't need much detail). This assignment can be turned in whenever you read the article. Submit the following:

1. The title of the scientific paper.
2. A link to the scientific paper.
3. A link to, or screenshot of, your retweet (link preferred).

Week 10: Tweet a sentence or two describing any scientific event—seminar, workshop, etc.—you have attended this semester and *not* previously tweeted about. Include the name of the event, who hosted it, and a link to their site (if applicable). This assignment can be turned in whenever you attend the event (you do not have to wait until Week 13). Examples include department seminars, etc. Feel free to use any scientific event, this is merely a suggestion. Submit the following:

1. The name of the event.
2. Who hosted the event.
3. The date of the event.
4. The website of the event (if applicable).
5. A link to, or screenshot of, your tweet (link preferred).

Week 11: Tweet a sentence or two about your semester (not necessarily this class). You will be graded equally for a positive or negative tweet as we are not trying to force good publicity for this lab. A tweet on your poster presentation from Week 13 is also acceptable. Submit the following:

1. A link to, or screenshot of, your tweet (link preferred).

**Appendix 4. Carbon Plate Calculations**

**The following is a guide to the components of the carbon substrate utilization experiment (weeks 2,4,6).**

**Sources tested in C-substrate experiments:** L-Arginine, L-Cysteine, L-Histidine, L-Isoleucine , L-Leucine, L-Lysine, L-Methionine, L-Phenylalanine, L-Threonine, L-Tryptophan, L-Tyrosine, L-Valine, L-Glutamine, Dextrose, Ribose, Pyruvate, Citrate, Oxaloacetic acid, Acetate, Succinate, a-ketoglutaric acid, Urea, Glycerol, Glycine, Choline chloride, Cyanate, DMSO, DMSP, B1/Thiamine, B2/Riboflavin, B7/Biotin, B9/Folic Acid, B12, Myo-inositol, Sucrose, Fructose, Glucose, L-Ornithine, Serine, and Glutamate

**Controls for C-Substrate experiments**: uninoculated carbonless medium, inoculated carbonless medium, inoculated isolation medium, uninoculated isolation medium

**Final concentration of carbon source needed in each well:** 0.5 µM
**Media volume in each well of plate:** 1.5 mL

**Directions:**
1. Make a stock solution of each carbon source in Milli-Q water.
2. Filter sterilize the stock with a 0.2 µm filter.
3. Store at 4˚C until use.
4. Use the equations below to determine the volume of carbon stock needed in each well to get 0.5 µM of carbon in 1.5 mL of medium per well.

**Calculations:**

1. Use the following equation to determine the mass of reagent needed to make a carbon stock at 0.00015M:

Mass = formula weight * Volume MilliQ (L) * concentration (M)

Therefore, if making a stock of Arginine with a formula weight of 210.7, the calculation is as follows:
Mass = 210.7 * 0.08 L * 0.00015 M = 0.00253 grams needed

2. Use the following equation to determine the volume of carbon stock needed for each well of the carbon plate:

Volume of carbon stock = (Concentration desired in well * Final volume in well) / Concentration of the carbon stock

Therefore, if the carbon stock is made at a concentration of 0.00015 M and we want to get a concentration of 0.5 µM (0.0000005 M) in our 1.5 mL wells, the calculation is as follows:

Volume to transfer into plate = (0.0000005 M * 1.5 mL) / 0.00015 M
Volume to transfer into plate = 0.005 mL = 5 µL

3. Use the following equation to determine the volume of culture needed to inoculate the carbon plate:

Volume to inoculate = (Concentration of culture desired in well * Final volume in well) / Concentration of culture

Therefore, if starting cell density is $5 \times 10^{6}$ cells·mL$^{-1}$ and we want to get a concentration of $1 \times 10^{4}$ cells·mL$^{-1}$ in our 1.5 mL wells, the calculation is as follows:

Volume to inoculate = ($1 \times 10^{4}$ cells·mL$^{-1}$ * 1.5 mL) / $5 \times 10^{6}$ cells·mL$^{-1}$

**Appendix 5. Writing Assignment 1: Carbon**

Use what you have learned about bacterial interactions with carbon for writing assignment #1. Format your writing as follows:

Introduction:

- Present information in a "funnel" that begins with the most general information and ends with the goal of the project
- General statement about the importance of bacteria
- Relate bacteria to carbon
- Why do we care about bacteria and carbon together?
- Why are we doing this experiment?
- Hypotheses about the experiment and controls

Methods:

- What did we do? List volumes and concentrations -be specific.
- Avoid including information about labeling or type of pipette used.

Results:

- What should the result table look like for the entire class? Create the table but leave it empty until all results are comepleted. .

Discussion:

- What controls did we place?
- What kind of results could we see with the controls?
- What would each of these mean?

**Rubric:**

| Section | Category | Points |
|---|---|---|
| Style | | 5 possible |
| | Name, Section Number, Section Headings, Double-spaced | 4 |
| | Title | 1 |
| Introduction | | 15 possible |
| | General funnel of information | 2 |
| | Define bacteria and their importance | 2 |
| | Importance of carbon to life | 3 |
| | Diversity of bacterial carbon usage and rationale for experiment | 3 |
| | Hypothesis | 3 |
| Methods | | 10 possible |
| | Sterile technique | 2 |
| | Volume of media, carbon source, culture | 5 |
| | Carbon substrate | 3 |
| Results | | 10 possible |
| | Table of carbon substrates with appropriate caption | 6 |
| | Written explanation of results | 4 |
| Discussion | | 10 possible |
| | Discussion of possible controls and results | 3 |
| | Support or reject hypothesis and why | 4 |
| Citations | | |
| | Copy-and-paste information in text | -6 per instance |
| | Invalid sources | -1 each instance |
| | No in-line citations | -3 each instance |
| | No citations | -5 |
| **Total** | | |

**Appendix 6. Presentation #1: Carbon Substrate Assignment**

Each person has been assigned a carbon source that we will test whether our organism can use as the sole carbon substrate for growth. This presentation will give you practice obtaining relevant primary literature online to answer the questions below and help you familiarize yourself with your substrate. Your instructor has assigned you to groups, and each person per group will prepare one slide about their carbon source with the following information:

1. The name of the carbon compound
2. The chemical formula of the compound
3. A picture of the compound's structure
4. A fact about where this compound is naturally found
5. A fact about how this compound is relevant to bacteria in the environment
6. Cite your source(s) in the footnote of your slide

You will each be given a maximum of two minutes to present your substrate. Be sure to practice this presentation to ensure concise wording and appropriate timing.

**Appendix 7. Homework #1: Downloading R and RStudio**

Download R:

1. Go to the following address: https://cran.cnr.berkeley.edu
2. Select the download that is most appropriate for you (Linux, Mac, or Windows)
3. Follow any download instructions that appear

Download RStudio:

1. Go to the following address: https://www.rstudio.com/products/rstudio/download/
2. Choose the free RStudio Desktop Open Source License option
3. Follow any download instructions that appear

Show your instructor the downloaded programs at the start of class.

**Appendix 8. Code for Growth Curves: Student Version**

```
# File: Growth_Curves.R
# Source: Lanclos et al., 2018
# Authors: V. Celeste Lanclos, Alex Hyer, Jordan Coelho

# Useful reminders:
# 1. Any text after "#" is ignored by R. Use this feature to take notes.
# 2. You need to change any text in ALL CAPS to match your data file.
# 3. RStudio let's you export graphs from the Plots tab. NO SCREENSHOTS!

# This script takes a CSV with the following columns as input:
#
# Salinity (int) or Temperature (int): the salinity or temperature at which the culture was grown
# Day (int) or Hours (int): how long the culture has been growing
# Replicate (int): trial number at a given salinity
# Cell.Count (int): the concentration of cells counted on each day

# Purpose:

#
rm(list=ls())

#
install.packages("ggplot2")

#
library(ggplot2)

#
DATA_NAME <- read.csv("NAME_OF_CSV_FILE_HERE.csv", header = T)

#
DATA_NAME$Replicate <- factor(DATA_NAME$Replicate, levels = unique(DATA_NAME$Replicate))

#
ggplot(DATA_NAME, aes(x = COLUMN_FROM_CSV_FOR_X_AXIS, y =
COLUMN_FROM_CSV_FOR_Y_AXIS, color = COLUMN_FROM_CSV_FOR_SERIES, fill =
COLUMN_FROM_CSV_FOR_SERIES)) +
  geom_line() +
  facet_wrap(~COLUMN_FROM_CSV_FOR_INDEPENDENT_VARIABLE) +
  labs(x = "X-AXIS_TITLE", y = "Y-AXIS_TITLE",
       title = "FIGURE_TITLE") +
  scale_y_log10() +
  theme_bw()
```

**Appendix 9. Writing Assignment 2: Temperature**

Use what you have learned about the effect of temperature on bacterial growth for writing assignment #2. Format your writing as follows:

Introduction

- -Funnel of information:
- -What is temperature, and why do we care about it in a bacterial sense?
- -Scope/diversity of temperatures that bacteria are found to thrive in
- -Why should we test temperature in our bacteria?
- -Hypothesis. Expected temperature range of the bacteria and why?

Methods

- -What did we do?
- -Be sure to include things such as sterile technique, media, media modifications if needed, temperatures, volumes, etc.

Results

- -What is the temperature min, max, and optimum for your organism
- -Graph – what we plotted in class.
- -Figure caption

Discussion

- -How does the data support or reject your hypothesis?
- -Is this temperature range surprising given the isolation locations?
- -What kind of data are our controls here?

**Rubric**

| <u>**Section**</u> | <u>**Category**</u> | <u>**Points**</u> |
|---|---|---|
| **Style** | | 5 possible |
| | Name, Section Number, Section Headings, Double-spaced, Title appropriate | 5 |
| **Introduction** | | 15 possible |
| | General funnel of information | 2 |
| | Define temperature and the relationship between bacteria and temperature | 5 |
| | Diversity of temperature for bacteria and rationale for experiment | 5 |
| | Hypothesis | 3 |
| **Methods** | | 10 possible |
| | Sterile technique | 1 |
| | Volume of media and culture | 3 |
| | Temperature range and replicates | 4 |
| | Description of how graphs were made | 2 |
| **Results** | | 10 possible |
| | Cardinal temperatures | 2 |
| | Temperature growth curve and figure caption | 6 |
| | Paragraph explaining results of graph | 2 |
| **Discussion** | | 10 possible |
| | Controls | 3 |
| | Support or reject hypothesis and why | 4 |
| | Temperature range in connection to the environment | 3 |
| **Citations** | | |
| | Copy-and-paste information in text or no citations | 0 on paper |
| | Invalid sources or no in-line citations | -3 each instance |
| **Total** | | **50** |

**Appendix 10. Homework #2: Growth Curves**

Now that you are all professionals at plotting temperature data in RStudio, your homework is to do the following:
1. Download the two .csv files containing temperature data from the other two sections' organisms
2. Input the data into RStudio
3. Change the R code to match headings or regroup data if needed
4. Change the title to the strain number and your name
5. Plot the data and save a PDF to submit

Write a short paragraph of what you think about the data that you see. This should include the differences and similarities of each organism's min, max, and optimum. Your final product will be two growth curve plots and one paragraph interpreting the data.

**Appendix 11. Homework #3: Poster critique**

Find a scientific poster displayed somewhere on campus or use one provided by your instructor to answer the following questions:

Based on the TITLE ONLY, what do you think the poster will be about?

What was the main research question and was the title an accurate description of it?

Was the main conclusion clearly stated? What was it?

Were the major ideas of each table/figure clearly stated in the table/figure legend? Were the tables/figures themselves clear and easy to understand? Were they all necessary? Please comment.

Was the order of sections in the poster formatted in a logical/easy to read way?

What did you like best about this poster?

What did you think was the worst thing about this poster?

How long did it take you to understand this poster?

Please feel free to add pages in order to fully explain your responses

**Appendix 12. Growth Rates Practice**

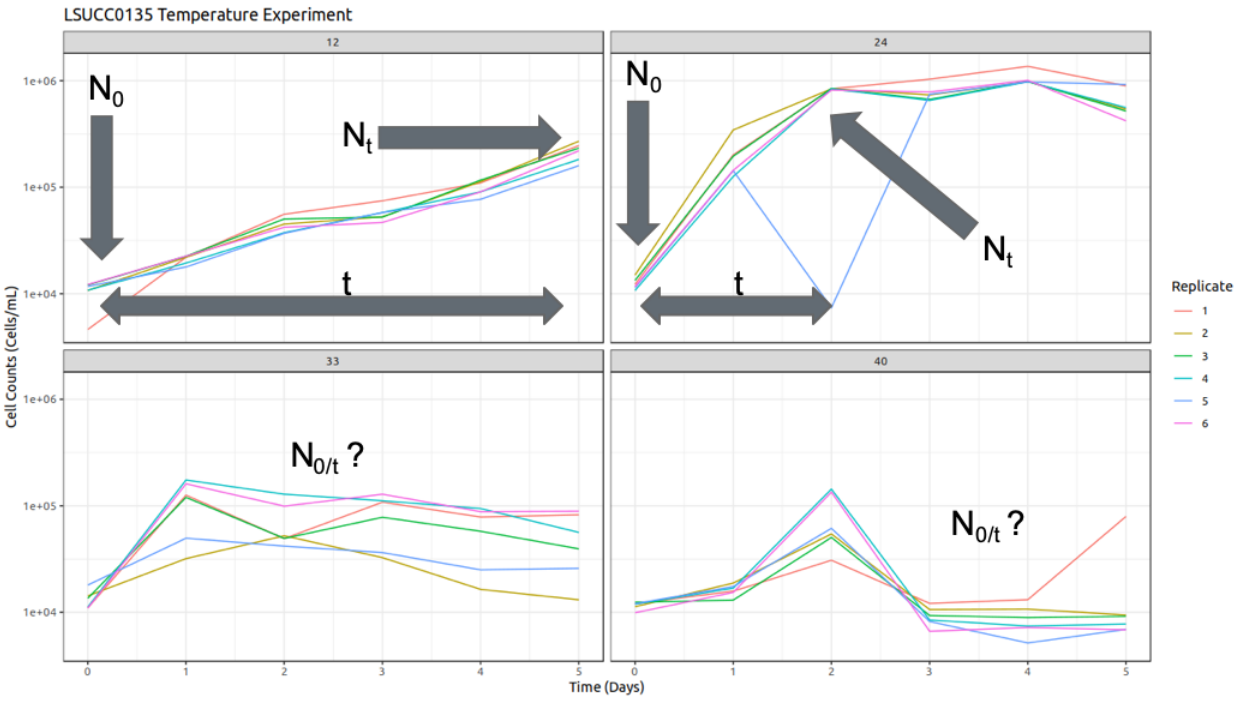

To find the growth rate of an organism, use the following:

Number of generations: $n = (\log(N_t) - \log(N_0)) / \log(2)$

number of generations = (log(#cells at end of log phase) - log(#cells at start of log phase)) / log(2)

0.301 = log(2)

Generation time: $g = n/t$

generation time = number of generations / elapsed time

Growth rate (k): $k = 1/g$

growth rate = 1 / generation time

---

If: $N_0$= 1,000 cells · $mL^{-1}$ $N_t$= 5,000 cells · $mL^{-1}$ t = 120 hours

Then:

n = (log(5,000) - log(1,000)) / 0.301 = 2.322 generations

g = 2.322 / 120 = 0.01935 generations per hour

0.01935 generations per hour * 24 hours/day = 0.4644 generations/day

k = 1 / 0.4644 = 2.153 days per generation

Interpretation: This organism doubles every 2.153 days under the given conditions.

**Appendix 13. Growth Rates: Student Version**

```
# File: Growth_Curves.R
# Source: Lanclos et al., 2018
# Authors: V. Celeste Lanclos, Alex Hyer, Jordan Coelho

# Useful reminders:
# 1. Any text after "#" is ignored by R. Use this feature to take notes.
# 2. You need to change any text in ALL CAPS to match your data file.
# 3. RStudio lets you export graphs from the Plots tab. NO SCREENSHOTS!

# This script takes a CSV with the following columns as input:
#
# Temperature (int) or Salinity (int): the temperature or salinity at which the culture was grown
# Replicate (int): trial number at a given temperature
# Rate (float): the rate at which a replicate grew

# Purpose:

#
rm(list=ls())

#
install.packages("ggplot2")

#
library(ggplot2)

#
DATA_NAME <- read.csv("NAME_OF_CSV_FILE_HERE.csv", header=T)

#
DATA_NAME$Replicate <- factor(DATA_NAME$Replicate, levels=unique(DATA_NAME$Replicate))

#
ggplot(DATA_NAME, aes(x = COLUMN_FROM_CSV_FOR_X_AXIS, y =
COLUMN_FROM_CSV_FOR_Y_AXIS)) +
  geom_jitter(size = 5, width = 0.0, height = 0.0, alpha = I(0.5)) +
  geom_smooth(method = "auto", formula = y ~ x, span = 0.85, se = FALSE) +
  labs(x = "X-AXIS_TITLE", y = "Y-AXIS_TITLE",
       title = "FIGURE_TITLE") +
  scale_x_continuous(breaks = c(CONDITION1, CONDITION2, CONDITION3, ETC),
                     labels = c("CONDITION1", "CONDITION2", "CONDITION3", "ETC")) +
  theme_bw()
```

**Appendix 14: Homework #4: Growth Rates**

After our in-class review of the new R code for plotting growth rates, plot growth rates for the temperature experiment. Submit a brief interpretation of results and a PDF of the curve with appropriate title, labels, and figure caption.

Your final product will be one plot of growth rates and a written interpretation of the data.

**Appendix 15. Final Writing**

For this writing assignment, you should stop thinking of carbon substrate usage, temperature, and salinity as unrelated experiments. Rather, remember that this entire course is actually a single project to characterize our organisms. This writing assignment is meant to be a formal, comprehensive scientific report about the organism that you have grown to know and love. This paper should be written so that an audience completely unrelated to our class could understand it. This assignment will be broken into multiple parts:

**Part 1**

Title: Should be an original descriptive title that includes key words of the project

Introduction:

-This section MUST include at least three outside references that you find on your own to support your ideas
-Why are bacteria important to study?
-Explain why carbon usage, temperature, and salinity are important factors relating to bacteria and link that information back to our isolation site.
-Introduce our organism. Include where it was isolated, how it was isolated, anything important about the media, why this media is good/bad, where the isolate currently exists, anything that had been published prior to this project, etc.
-Introduce our research questions, hypotheses, etc.

Methods:

-Be comprehensive, nonrepetitive, and concise.
-Use separate subheadings for:
    -Sterile Technique
    -Cell Counts
    -Carbon substrate usage
    -Temperature
    -Salinity

References:

-This should have a minimum of 5 references that are listed here and also have in-text citations throughout the paper.

**Part 2:**
Use what you have turned in for Final Writing Part 1 and add in the following sections:

Methods:
- Add in information for plotting data in RStudio.

Results: At this point, results should be comprehensive!
You should have the following figures with captions:
  - Carbon Table
  - Temperature Growth Curves
  - Temperature Growth Rates
  - Salinity Growth Curves
  - Salinity Growth Rates
- Data in figures must be also typed out and reported in the text. Reference this in the text as relevant.
  - EX: LSUCC0135 is a curved rod (Figure X).
- Break this category into relevant subsections as needed.
- There should be no interpretation of these results.

Discussion:
- Use this section to tie in our results to what we know from existing literature and things that we have learned in class. This section is for the bigger picture connections from what we see in our data to existing data and the world around us.
- There should be citations here

Future Directions:
- What would you like to see happen next in this series of experiments with our organism?

References:
- Follow APA style and list in order of appearance
- Use numerical in-text citations

**Part 3: Peer Edits**

Come to class on with your entire Final Writing printed double spaced with NO NAME. We will be doing peer-review in class, and these should remain anonymous to your classmates. Choose a codename that will not identify you and print it clearly on every page of the document. When we get to class, I will document who belongs to each codename to assign grades. Part of your grade for this final writing assignment comes from how well you peer review for your classmates. There is no way to make up these peer review points. You must come to class. The peer review is worth 40 points towards the 200 points of this assignment.

**Rubric:**

| **Section** | **Category** | **Points** |
|---|---|---|
| **Peer Review** | | 40 possible |
| | Thoughtful comments and relevant feedback on peer’s paper | |
| **Style** | | 10 possible |
| | Name, section number, appropriate title, section headers, appropriate grammar, easy to read and concise | 7 |
| | Unit symbols where necessary | 3 |
| **Introduction** | | 40 possible |
| | Importance of bacteria | 5 |
| | Synthesis of bacterial relationship with carbon, temperature, and salinity | 15 |
| | Introduce organism and any known information about it | 7 |
| | Describe the rationale of this experiment and brief description of it | 8 |
| | Hypothesis | 5 |
| **Methods** | | 30 possible |
| | Sterile Technique | 3 |
| | Counting: Instrument, stain, limit of detection | 4 |
| | Carbon | 4 |
| | Temperature | 4 |
| | Salinity | 7 |
| | Computation: R or RStudio | 4 |
| | Computation: ggplot2 | 4 |
| **Results** | | 30 possible |
| | Carbon Table and caption | 5 |
| | Temperature growth curves and caption | 5 |
| | Temperature growth rates and caption | 5 |

| | | |
|---|---|---|
| | Salinity growth curves and caption | 5 |
| | Salinity growth rates and caption | 5 |
| | Written results in addition to the figures | 5 |
| **Discussion** | | 30 possible |
| | Connection to ecology/ bigger idea | 20 |
| | Well thought-out acceptance/rejection of hypothesis | 10 |
| **Future Directions** | | 15 possible |
| | Thoughtful idea of a follow-up experiment | |
| **References** | | 5 possible |
| | In-text citations and properly formatted citations | 5 |
| **Total** | | **200** |

**Appendix 16: Poster Assignment**

Now that you've written your final paper, let's practice another form of science communication. A poster is a great way to discuss your research and present your data, while personally connecting with your audience. The general construct of a scientific poster is similar to the scientific papers you have been writing; however, the poster is concise and highlights the key points and findings. Essentially, condense your paper into poster format.

Like we discussed in class, your poster should be aesthetically pleasing and not cluttered.

Your poster needs to include:

1) Title
2) Your name and your lab partner's name, and the department where the research was conducted
3) Introduction
    a. Briefly highlight the relevant background, what makes this research important/what gap in the knowledge we are filling with our work, and your research question/hypothesis.
4) Methods
    a. Briefly discuss how we conducted our research and make sure to explain each experiment distinctly. Feel free to be creative with figures, charts, diagrams, etc
5) Results
    a. Provide the data produced from each experiment
    b. Make sure that each figure/table has an informative caption
6) Discussion
    a. Briefly discuss the main findings of our investigation and any conclusions we can draw from them
    b. Briefly discuss the relevance of these conclusions to the broader scientific community
    c. Bullet points are encouraged here
7) Acknowledgements
    a. The purpose of this section is to thank anyone who made this research opportunity possible
    b. Be sure to thank the department, the university CURE program and any staff that coordinate/direct the program, the instructor of record, the Principle Investigator, and your Laboratory Teaching Assistant.
8) References
    a. APA format and numerical in-text citations

Make sure your poster is sized for:

Width: 48 inches
Height: 36 inches

For the presentation, aim for 3-5 minutes.

Things to keep in mind:

-Your poster should be easy for your reader/audience to read and follow along
-No clutter, no distracting background
-Make sure the font is large enough/figures are readable
-Keep your poster spatially organized

**Rubric**

| **Section** | **Category** | **Points** |
|---|---|---|
| Design and Layout | | |
| | Organization | 10 possible |
| | Readable figures and text | 10 possible |
| Heading | | |
|  | Names and Affiliation | 5 possible |
| | Clear and accurate title | 5 possible |
| Content | | |
| | Introduction | 20 possible |
| | Methods | 10 possible |
| | Results | 10 possible |
| | Figures, Tables, and Captions | 20 possible |
| | Discussion | 20 possible |
| | Acknowledgements and References | 10 possible |
| Presentation | | |
| | Time | 10 possible |
| | Professionalism and body language | 20 possible |
| **Total possible** | | 150 possible |

**Appendix 17. Elevator Pitch Assignment: Modified from Becky Carmichael, Kyle Sirovy, Scott Kosiba, Mindy Brooks, and Courtnie DiCapo.**

Part 1: In class assignment

Scientists communicate their findings through a number of mediums to a wide variety of audiences that range from fellow scientists within the same field of study to the general public. The composition of your audience should always dictate how you discuss your science. If you were to present the findings of your research to a room of non-scientists but your presentation was more appropriate for fellow scientists, the presentation would fall flat. What's the point of communicating if those receiving that communication don't understand ?

The goal of this assignment is to expose you to one of the many ways scientists communicate their research to specific, well-defined audiences and highlight the broader impacts of their research in a way that is both relevant and interesting to that audience. This exercise will help you identify your audience, cater your communications with that audience, and aid in the development of an elevator pitch for your final posters.

Please read the following questions before listening to the science communication. Feel free to make notes throughout and then provide detailed responses to these questions:

Title of science communication:____________________________________________________

Presenter's name, title, and affiliation:________________________________________________

Type of science communication (podcast, seminar talk, blog post, etc.):________________________

1. Do you feel that the presenter has a strong grasp of the research they conduct? What gave you this impression? What aspects could you model for your own pitch/what do you want to incorporate?

2. Who is the target audience for this communication? How can you tell?

3. Were they effective at communicating their research to this audience? What did you find specifically effective or not? How did the presenter "hook" the listener and sustain attention?

4. List the main takeaways (broader impacts/results) of the research.

5. Was the importance of their research clear? What was it?

6.  What did you like best about the presentation of the material?

7. What clarifying or engaging question would you want to ask the presenter? What was left unanswered?

8. After listening to the science communication, what aspects of the presenter's delivery style will you model for your own presentation?

9. List specific aspects from your CURE research that you will highlight including the big picture take-away, and relatable example.

Part 2: Homework

An elevator pitch is a short talk used to introduce yourself and your research to others and includes the question being addressed, the importance of the work, and major findings. You've already evaluated a science communication piece (podcast episode, seminar talk, published paper, blog post, etc.) and analyzed that researcher's pitch. Now, you will use your previous evaluation to craft an elevator pitch on existing research. Use the instructor suggestions to find a relevant piece of scientific work and create an elevator pitch on that work. Fill out the prompts below:

Title of scientific research:________________________________________________

Presenter name(s), title(s), and affiliation:________________________________________

Type of science communication (podcast, seminar talk, blog post, etc.): ______________________

When composing any sort of science communication, be it a poster or paper to be submitted to a peer-reviewed journal, two components are essential: A well-defined target audience and a clear story.

*Step 1. Identify your target audience*
The reason many communications of science fail to resonate with those listening or reading is because the authors did not adequately identify who they are communicating to. By zeroing in on a specific audience you can cater components of your speech to their level of interest, understanding of complex relationships and methodologies, and formulate broad impacts that will resonate.

1. What are some aspects of your target audience that you can define? Think: education level, location/affiliations(s), occupation(s), age, familiarity with broad topics you'll discuss. List as many specifics as possible. Example: *College educated, mostly PhD and MS degree seeking.*

Your target audience:

2. Using your response above, create a single person that fits into this target audience. In a few sentences, write a general description of this person—this is who you'll be crafting your pitch for!

*Step 2. Find your story*
When communicating science in any form, it is crucial to have a clear idea of the story you want to tell. A clear story makes communicating your results more interesting to your audience and helps to convey your desired message effectively.

3. Write a two-sentence summary of the research question and the answer (main result/conclusion) to that question. This two-sentence summary will form the backbone of your elevator pitch.

4. From the two-sentence summary you’ve created above, you should include a brief summary of the approach used to obtain the main result. This does not need to be overly detailed but should be clear and specific enough that your audience understands the general approach taken. Now, rewrite your two-sentence summary to include a sentence (or two) about the general approach taken to obtain the main result--this should be sandwiched between your research question and the main result/conclusion.

5. Consider why the main result is important from the perspective of your audience. It’s not enough to state a result if nobody understands the point of it all. Rewrite your two-sentence summary on the back of this sheet to include a sentence (or two) discussing why the results are important from an ecological or biological perspective and also why this should be interesting/relevant to your audience.

6. Practice saying your elevator pitch and come to class ready to present it to your classmates in small groups.

**Appendix 18. Informal Essay #2**

In paragraph format, write an informal essay that highlights your experience now that you have taken part in a Course-based Undergraduate Research Experience (CURE) lab. There is no strict word count on this assignment, but your goal should be about half a page.

Here are some topics to cover:

-After revisiting your Blog Post #1, address your overall thoughts about your experience in the class. Was your initial impression with the course consistent throughout the semester? Did your views on research and what it means to be a researcher shift throughout the semester? Would you now consider yourself a researcher or at least capable of being a researcher?

-Has your confidence in your lab abilities grown this semester? Do you feel that any skills you have learned are relevant to you and your goals? Would you recommend a CURE lab to a friend?

**Appendix 19. Example Final Exam Fall 2018**

Experimental Theory (150 points)
General:

1. What were 2 of the ways that we used sterile technique to minimize possible contamination in this lab? (2 points each, 4 points total)

    1)

    2)

2. Describe a specific way that bacteria uniquely affect their environment. (4 points)

3. List 3 reasons that bacteria can be beneficial to humans. (2 points each, 6 points total)

    1.

    2.

    3.

4. What are the two types of controls that we should have with experiments? (2 points each, 4 points total )

5. Define the predicted outcome of each of the controls from above: (2 points each, 4 points total)

    1.

    2.

6. Which of the following nutrient cycling do bacteria NOT help with: (2 points)
    a. carbon
    b. sulfur
    c. nitrogen
    d. phosphorous
    e. bacteria help with all of the above
    f. bacteria don't participate in any of the above

Scientific literature:

7. What is the difference between a motivation and an objective? (2 points each, 4 points total)

    Motivation:

Objective:

8. What are the 6 major sections of a scientific paper? (1 point each, 6 points total)

9. Fill in the section of a scientific paper that the following information would be found: (2 points each blank, 8 points total)

| *Excerpt from paper* | *Section of paper* |
|---|---|
| Within the environment, chromium mainly persists in two forms: Cr(III) and Cr(VI) (Bartlett, 1991). Cr(VI) is highly toxic, soluble, and can be easily transported across cell membranes of both eukaryotic and prokaryotic organisms via sulfate and other active transporters (Ackerley et al., 2004b; Cheng, Holman & Lin, 2012). | |
| Though a "core" metabolic and genomic structure was seen among our isolates, our data suggests that Cr(VI) reduction discrepancies within these isolates could be related to strain-level genetic and metabolic variation. Further, chromate resistance may be intertwined with the ability of a bacterium to reduce and transport chromate as well as the type of stress response the organism might have. | |
| Each genome did contain genes with sequence homology to the chromate reductases, chrR and yieF, of non-model organisms. The putative chrR-like genes found in the isolates are homologous to a chrR gene (GenBank accession number AM902709) found in T. scotoductus. (Opperman, Piater & Van Heerden, 2008). | |
| Whole genome shotgun sequencing was performed by multiplexing the genomic DNA onto one lane using the Illumina HiSeq 2000 platform with 100 bp paired end reads using V2 chemistry at Cincinnati Children's Hospital Medical Center's Genetic Variation and Gene Discovery Core Facility. | |

Carbon:

10. Autotrophs are associated with organic / inorganic carbon. (2 points)

11. Heterotrophs are associated with organic / inorganic carbon. (2 points)

12. Rocks, respiration, and sediment are sources of organic / inorganic carbon in aquatic ecosystems. (2 points)

13. In 3 sentences or less, why did we perform transfers when characterizing the carbon usage of your organism? (3 points)

Use the following result table to answer questions 14-17:

| Well | Carbon Source | P1 | P2 | P3 |
|---|---|---|---|---|
| A1. | Sodium acetate | + | - | - |
| A2. | Sodium succinate | + | - | - |
| A3. | Sucrose | + | - | - |
| A4. | Urea | + | + | + |
| A5. | Glycine | + | + | + |
| A6. | Bacteria + media | + | + | + |
| A7. | Bacteria + carbonless media | + | + | + |
| A8. | Carbonless media only | + | + | + |

14. Which wells contain the carbon sources that this organism can use? (3 points)

15. Which wells contain the controls? Label them as positive or negative controls. (6 points)

16. Can we trust this data? (2 points)

17. Why or why not? (3 points)

18. Fill in the blank to go through the protocol for the carbon experiment: (3 points each blank, 15 points total)

We started the first carbon plate with 1.5mL of____________________media already in the wells of Plate #1. We then added our ____________________to give our organism food to grow. Then we inoculated Plate #1 with our bacteria- ____________________._We waited for 2 weeks, then Plate #1 was counted using in the Lab. The source of the bacteria for Plate #2 was ______________________ since this portion of the experiment was a transfer rather than an inoculation.

19. What is NOT one of the fates of carbon for a heterotrophic bacterium? (3 points)

a. CO2

b. Inorganic Carbon

c. Waste

d. Biomass

20. T/F: All bacteria found in the same environment use the same types of carbon substrates. (1 point)

Temperature:

21. T/F: Bacteria can regulate their temperature. (1 point)

22. Rank from coldest to hottest: psychrophile, thermophile, hyperthermophile, hyper-psychrophile, mesophile (4 points)

23. Since temperature directly affects bacterial growth rate, if a bacterium is growing in a system that is outside its optimum temperature range it will be more / less active. (2 point)

Use the following graph from XX to answer questions 24-27:

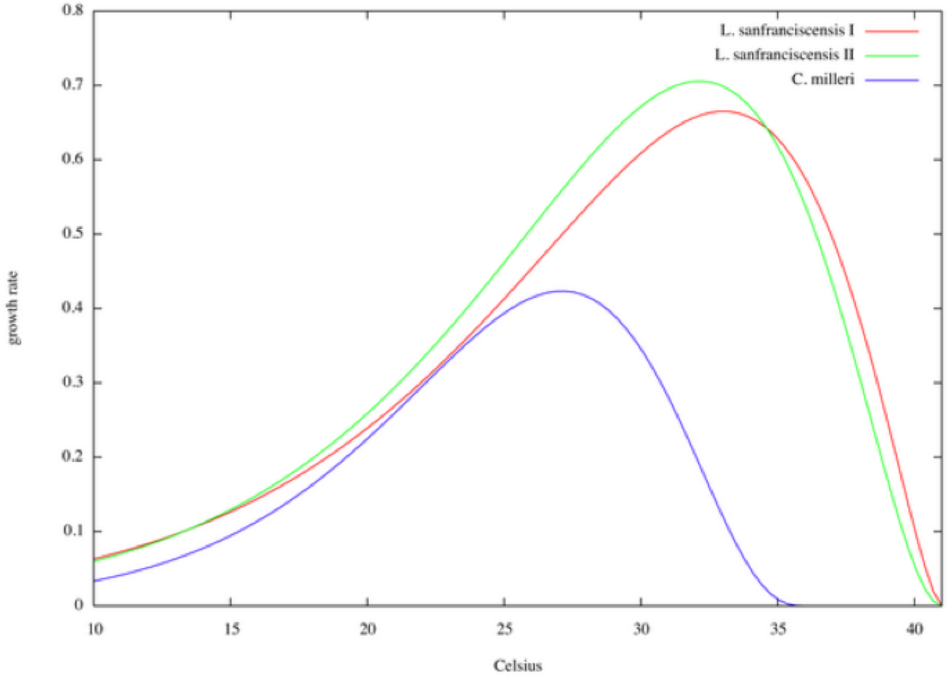

24. Which organism has the lowest range of possible temperatures? (3 points)
    a. L. sansranciscensis I
    b. L. sansranciscensis II
    c. C. milleri

25. If all of these organisms were in a flask together, which do you think would dominate the culture? Hint: Which would outgrow the others? (3 points)
    a. L. sansranciscensis I
    b. L. sansranciscensis II
    c. C. milleri

26. What temperature range would you expect to see all of these organisms in the environment co-existing? (3 points)

27. Explain your answer for number 26 above. What about the graph tells us that this is the correct range? (4 points)

Salinity:

28. T/F: In the salinity experiment, we used media with different types of salts to figure out which salts the organism liked best. (2 points)

29. T/F: If we adjust an organism's media from a salinity value of 5 to a salinity value of 10, it will definitely be okay because both of these values are in the brackish range. (2 points)

30. Osmosis is active/passive. (1 point)

31. Osmolyte transport is active/passive. (1 point)

32. T/F: Defining salinity tolerances helps in understanding bacterial distribution. (1 point)

33. If a cell is placed from a fresh habitat into a salty habitat, describe what happens to the water inside of the cell. Draw an image if you'd like. (5 points)

34. Why are coastal areas commonly brackish? (5 points)

35. The saltiest type of water is: (2 points)
- a) brackish
- b) briny
- c) fresh
- d) saline
- e) super saline

Microbial Growth and Growth Rates:

36. Fill in the blanks or circle the appropriate answer pertaining to microbial growth and growth rates. (3 points each, 15 points total)

Bacteria reproduce through binary___________________. This process is when one cell becomes two through (sexual / asexual reproduction). Because of this type of reproduction, cultures of bacteria experience______________________ growth, which is indicated in the growth curves as the part of the curve with the steepest slope. If we can find the number of cells at the inflection points of a growth curve, we can calculate the ______________________ of the culture. This value allows researchers to easily compare microbial growth. Once plotted into a curve, if forms a (sigmoidal / bell curve) shape in which we can see the minimum, maximum, and optimum condition of a bacterial culture.

37. T/F: The way an organism will grow in a lab is the exact way that it will grow in the environment (2 points)

38. T/F: Generation time is the time it takes for one cell to become 2. (1 point)

39. The equation: n = (log(Nt) - log(N0)) / 0.301 is used to calculate what? (3 points)

a.The time at which lag phase ends
b.The time between Nt and N0
c. The cell count at the beginning of stationary
d. Generations between Nt and N0
e. Generations per time
f. Time per generation
g. Growth rate

40. The equation: g = n/t is used to calculate what? (3 points)

a.The time at which lag phase ends
b.The time between Nt and N0
c. The cell count at the beginning of stationary
d. Generations between Nt and N0
e. Generations per time
f. Time per generation
g. Growth rate

41. The equation: k= 1/g is used to calculate what? (3 points)

a.The time at which lag phase ends
b.The time between Nt and N0
c. The cell count at the beginning of stationary
d. Generations between Nt and N0
e. Generations per time
f. Time per generation
g. Growth rate

Bonus: Explain in detail something that you learned this semester that was not on the exam. You can earn up to 5pts.

**Appendix 20. Poster Symposium Worksheet**

| **Criteria** | Poster #:________ |
|---|---|
| **1. Content:** short, understandable | 1 2 3 4 5 6 7 8 9 10 |
| **2. Organization/clarity:** easy to follow and understand, flows well | 1 2 3 4 5 6 7 8 9 10 |
| **3. Audience:** detail appropriate; big picture catered to audience | 1 2 3 4 5 6 7 8 9 10 |
| **4. Completeness:** detail and depth is appropriate | 1 2 3 4 5 6 7 8 9 10 |
| **5. Volume:** Projects voice, appropriate for group size | 1 2 3 4 5 6 7 8 9 10 |
| **6. Pace:** Relaxed pace, easily understood given accent/vocal style | 1 2 3 4 5 6 7 8 9 10 |
| **7. Diction:** pronunciation is clear and deliberate | 1 2 3 4 5 6 7 8 9 10 |
| **8. Enthusiasm/energy:** shows interest through tone and energy | 1 2 3 4 5 6 7 8 9 10 |
| **9. Body language/posture:** does not fidget, upright posture | 1 2 3 4 5 6 7 8 9 10 |
| **10. Eye contact:** maintains eye contact, engages audience member | 1 2 3 4 5 6 7 8 9 10 |
| **11. Audience questions:** clear and thoughtful response | 1 2 3 4 5 6 7 8 9 10 |
| **OVERALL POSTER SCORE** | |

1. Based on your overall impression of the presentation, do you feel that they have a strong grasp of the experiment they conducted and its relevance to the field? Were they able to effectively communicate this to their **audience**? Be specific with your feedback.

2. What did you like best about the presentation of this poster?

3. What could have been improved or refined? Use the scores you provided to guide your response.

4. What clarifying or engaging question did you ask and what was the answer given? Did you feel that they answered it thoroughly and with the audience in mind?

**Appendix 21. Flow cytometry parameters (Modified from Bakshi et al. 2019)**

This protocol assumes that users have received the proper training in flow cytometry and understand how to use their equipment. The following parameters are used with the Guava easyCyte 5HT (Millipore) flow cytometer for enumeration:

**Gain settings**
Forward scatter 1
Side scatter 2.83
Green fluorescence 4.56
Yellow fluorescence 8
Red fluorescence 8

**Counting**
3,000 events or 90s.

**Controls**
Negative: unstained sterile medium
Negative: stained sterile medium
Positive: medium with a common marine bacterial heterotroph (e.g., LSUCC0096).

Additional details and gating examples can be found in:

Thrash, J. Cameron, Jessica Lee Weckhorst, and David M. Pitre. (2015) *Cultivating Fastidious Microbes.* In Hydrocarbon and Lipid Microbiology Protocols, vol. 4 (*Cultivation*). Edited by Terry J. McGenity, Kenneth N. Timmis and Balbina Nogales.

**Appendix 22. Dishwashing protocol**

The following steps should be used to ensure clean dishware:
-Rinse with copious amounts of water
-Scrub using brushes or sponge with soap and hot water
-Rinse with hot water
-Rinse with cold water
-Rinse with DI water
-Allow to air dry on racks

If the dishware has an open top ro will be used for measurement, cover with foil and put away.

Flasks and glassware should be acid washed overnight in 10% HCl. After acid washing:
-Rinse 4x with DI water
-Rinse 4x with nanopure water
-Allow to air dry on racks
-Cap flasks and bottles before autoclaving

**Appendix 23. R Code for Growth Curves: Instructor Version**

```
#' File: Growth_Curves_Instructor.R
#' Source: Lanclos et al., 2018
#' Authors: V. Celeste Lanclos, Alex Hyer, Jordan Coelho

#' This script graphs the growth curve of an organism at different salinities or temperatures.
#' The code below is in terms of salinity, but can easily be modified for temperature.

#' This is the Instructor Edition of this code and thus has the
#' "right answers" as well as useful tips. Please provide students with the
#' Student Edition and take time to note what information they need to
#' fill in for themselves.

#' This script takes a CSV with the following columns as input:
#'
#' Salinity (int) or Temperature (int): the salinity or temperature at which the culture was grown
#' Day (int) or Hours (int): how long the culture has been growing
#' Replicate (int): trial number at a given salinity
#' Cell.Count (int): the number of cells counted on each day
#'
#' You can, of course, edit the code and file to match your needs.

#' Removes any data from R's memory.
#' Note that this removes all environmental variables, please exercise caution.
#' In particular, don't execute this line after opening up an R Project file.
#' We strongly recommend that you execute this file in it's own R session.
#' This line is included to ensure a clean environment and prevents students
#' from unwittingly using previously stored data from their global environment
#' and getting perplexing results.
rm(list=ls())

#' Install the graphing software "ggplot2".
#' Comment this out/delete it if you've already installed it.
install.packages("ggplot2")

#' Load ggplot2 into R so that we can make our plot later.
#' This library will be removed by the `rm(list=ls())` which means students
#' can't proceed with plotting until they re-run this. It serves as a useful
#' check that they actually went through the whole file.
library(ggplot2)

#' Read your CSV file into R and saves the dataframe as "salinity" or "temperature".
#' Put the path to your file in between the quotation marks below.
#' You can use the tab key to more easily find your file in RStudio.
#' We found the GUI window was useful for students new to software.
salinity <- read.csv("salinity_growth_curve_example_data.csv", header = T)

#' Tells R to treat your replicates as categories instead of a number series.
#' We found that student's frequently skipped this line whenever they were
#' troubleshooting or trying to execute the code on their own for homework.
#' If this line is skipped, R will treat the replicates as a number series
#' which results in three obvious and incorrect features:
#' 1. There will only be a single series on their graph.
```

```
#' 2. The line is largely blue with black segments instead of a solid color.
#' 3. Each salinity on the x-axis will have a vertical spike.
#' If you notice these features on a student's graph, they have skipped this
#' critical line.
salinity$Replicate <- factor(salinity$Replicate, levels = unique(salinity$Replicate))

#' Render the graph using ggplot2.
#' Commenting out lines below and re-making the graph will help students
#' learn what each line does.
ggplot(salinity, aes(x = Day, y = Cell.Counts, color = Replicate, fill = Replicate)) +  # aes = aesthetic
  geom_line() +  # Make this graph a line graph
  facet_wrap(~Salinity) +  # Make a new graph for each salinity
  labs(x = "Time (Days)", y = "Cell Concentration (cells · mL^-1)",  # Change the graph labels
       title = "Test Organism Salinity Curve") +  # Change the graph title
  scale_y_log10() +  # Scale the y-axis by log10
  theme_bw()  # Change the theme to a clean black and white theme

#' Students can now export the graph using the Export function in RStudio.
#' Don't let students submit screenshots.
```

**Appendix 24. R Code for Rates: Instructor Version**

```r
#' File: Growth_Rates_Instructor.R
#' Source: Lanclos et al., 2018
#' Authors: V. Celeste Lanclos, Alex Hyer, Jordan Coelho

#' This script graphs the growth rate of an organism at different salinities or temperatures.
#' Note: The code below is for salinity but can be easily modified for temperature.

#' This is the Instructor Edition of this code and thus has the
#' "right answers" as well as useful tips. Please provide students with the
#' Student Edition and take time to note what information they need to fill
#' fill in for themselves.

#' This script takes a CSV with the following columns as input:
#'
#' Salinity (int) or Temperature (int): the salinity or temperature at which the culture was grown
#' Replicate (int): trial number at a given salinity or temperature
#' Rate (float): the rate at which a replicate grew
#'
#' You can, of course, edit the code and file to match your needs.

#' Removes any data from R's memory.
#' Note that this removes all environmental variables, please exercise caution.
#' In particular, don't execute this line after opening up an R Project file.
#' We strongly recommend that you execute this file in it's own R session.
#' This line is included to ensure a clean environment and prevents students
#' from unwittingly using previously stored data from their global environment
#' and getting perplexing results.
rm(list=ls())

#' Install the graphing software "ggplot2".
#' Comment this out/delete it if you've already installed it.
install.packages("ggplot2")

#' Load ggplot2 into R so that we can make our plot later.
#' This library will be removed by the `rm(list=ls())` which means students
#' can't proceed with plotting until they re-run this. It serves as a useful
#' check that they actually went through the whole file.
library(ggplot2)

#' Read your CSV file into R and save it as "salinity" or "temperature".
#' Put the path to your file in between the quotation marks below.
#' You can use the tab key to more easily find your file in RStudio.
#' We found the GUI window was useful for students new to software.
salinity <- read.csv("salinity_growth_rate_example_data.csv", header=T)

#' Tells R to treat your replicates as categories instead of a number series.
#' We found that student's frequently skipped this line whenever they were
#' troubleshooting or trying to execute the code on their own for homework.
#' If this line is skipped, R will treat the replicates as a number series
#' which results in three obvious and incorrect features:
#' 1. There will only be a single series on their graph.
#' 2. The line is largely blue with black segments instead of a solid color.
```

```
#' 3. Each salinity on the x-axis will have a vertical spike.
#' If you notice these features on a student's graph, they have skipped this
#' critical line.
salinity$Replicate <- factor(salinity$Replicate, levels=unique(salinity$Replicate))

#' Render the graph using ggplot2.
#' Commenting out lines below and re-making the graph will help students
#' learn what each line does.
ggplot(salinity, aes(x = Salinity, y = Rates)) +  # aes = aesthetic
  geom_jitter(size = 5, width = 0.0, height = 0.0, alpha = I(0.5)) +  # Graph data points
  geom_smooth(method = "auto", formula = y ~ x, span = 0.85, se = FALSE) +  # Draw average rate line
  labs(x = "Salinity", y = "Growth Rate (k)",    # Change the graph labels
       title = "Test Organism Salinity Growth Rate") +    # Change the graph title
  scale_x_continuous(breaks = c(6, 12, 23, 35),  # Manually set axis breaks to match salinity
                     labels = c("6", "12", "23", "35")) +  # Manually label the breaks
  theme_bw()  # Change the theme to a clean black and white theme

#' Students can now export the graph using the Export function in RStudio.
#' Don't let students submit screenshots.
```

**Appendix 25. Quizzes**

**Pipettes and Carbon**

1). You have three mechanical pipettes available for use: P10, P100, P1000.

A) Which is/are the pipette(s) that can hold 20 µL?

B) Which is/are the pipette(s) that can hold 100 µL?

C) Fill the boxes (right) to read as the volume readout should on the P10 pipette if you were to use it to deliver 9.2 µL volume.

2) List 2 reasons that all organisms need carbon.

3) What are the two general forms of carbon?

4) Explain the difference between autotroph and heterotroph:

5) Why do we care to test Carbon use in our isolates?

6) Which part of the Tree of Life are the "evolutionary winners"

Bacteria Archaea Eukaryotes

7) Are all bacteria bad for you? Explain something beneficial that bacteria can do. Be as specific as possible and use the back of the page if needed.

**Temperature**

1. The definition of temperature that we are using for this class is: “A measurement of the

_____________ _______________ of the molecules within a _____________.

2. Rapid movement of the molecules equates to (hot / cold) temperature.

3. List 2/3 effects temperature has on bacteria’s ability to survive:

   1.

   2.

4. T/F: Bacteria can regulate their own internal temperatures.

5. Why are temperature graphs shaped like a bell curve?:

6. Draw a diagram with the 3 Cardinal Temperatures. The y axis should be Growth Rate and the x axis should be Temperature:

7. BONUS: Draw the graph from above with the bell curves and labels of the 4 temperature -phile types we saw in class.

**Salinity**

1. Which is not a category to describe the salinity of water?
   a. Brackish
   b. Fresh
   c. hyperbrackish
   d. Briny
   e. saline

2. Order the classifications from Q1 from most fresh to most salty:

3. Osmosis is the active/passive movement of water through cell membranes in response to solute concentrations outside of the cell.

4. If solute concentration outside of the cell is higher than inside the cell, water:
   a. Moves inside the cell
   b. Moves outside the cell
   c. Moves in and out at an equal rate
   d. No movement

5. If solute concentration inside of the cell is higher than outside the cell, water:
   a. Moves inside the cell
   b. Moves outside the cell
   c. Moves in and out at an equal rate
   d. No movement

6. What is an osmolyte?

7. The following questions are in regard to our artificial seawater media that we use in class:

   a. Which paper was the one to create this media?

   b. What is the difference in medias that we are using for this salinity experiment? Your answer should go beyond they are different salinities and explore what makes them different salinities.

**Bacterial Growth**

1. Bacteria reproduce sexually/asexually.

2. Culture growth refers to what?

3. What is the technique we use to measure growth of our cultures?
    a. Colony forming units
    b. Flow cell cytometry
    c. Hemacytometer
    d. Optical density

4. Draw a standard bacterial growth curve, and include the phases of growth:

5. SYBR-green stains what part of the bacteria?
    a. Cell wall
    b. Mitochondria
    c. Proteins
    d. DNA

6. How do you add a comment in R Studio?

7. This is the code that we have been using. Circle the part of the code that allows you to change the axis labels. Put a square around the part of the code that allows you to change the way the data is grouped.

```
ggplot(Temp, aes(x =Day, y = Cell.Count, color= Replicate,  fill = Replicate)) +

    geom_line() +

    labs(x= "Time (Days)", y = "Cell Counts (cells · mL-1)",

    title = "LSUCC0117 Temperature Experiment") +

     theme_bw()+

     facet_wrap(~Temperature) +

     scale_y_log10()
```

8. What was the motivation of Henson et al. 2016?

9. What was the goal of Henson et al. 2016?

10. What was the measurement of cultivation success/failure in Henson et al. 2016?

**Growth Rates**

1. Define what a "growth rate" is.

2. Where do we get data to calculate growth rates?

3. What is generation time?
   a. Time it takes for cells to double in size
   b. Time it takes for a single cell to divide
   c. Growth of cells over a period of time
   d. Time it takes for the population size to double

4. What do we NOT need to calculate generation time?
   a. No
   b. k
   c. Nt
   d. n

5. Fill in the equation
   n= (log( )−log( )) / 0.301

6. Which is the line that assigns data for your x-axis and y-axis?

```
1.ggplot(nameyourdata, aes(x=XXXX, y=XXXX)) +
2. geom_jitter(size = 5, width = 0.0, height = 0.0, alpha = I(0.5)) +
3. geom_smooth(method = "auto", formula = y ~ x, span = 0.85, se = FALSE)
4. labs(x="Enter a label", y="Enter a label", title="Enter a label") +
5. scale_x_continuous(breaks=c(12,24,33,40),
6. labels=c("12","24","33","40")) +
7. theme_bw()
```

7. Is the code above for growth curves or rates?

8. What is the difference in axes for growth curves vs growth rates?

9. Draw what a growth curve graph looks like on the left and growth rate graph on right.

**Appendix 26: Example informal essay submissions**

**Student #1 Essay #1**-"I am an animal science, pre-vet major. My goal is to eventually earn my DVM and a PhD in either small animal soft tissue surgeries or exotics. I am very interested in research, particularly in studies that better our understanding of animals…I haven't heard much about CURE labs other than the fact that they are a great way to get your feet wet in undergraduate research. My only worry for this course is that I won't be able to keep up. However, I know I will try my best to be as successful as possible. I am very excited to be a part of a real research lab making new discoveries instead of doing "cookie cutter" labs and getting the same results as thousands of students before me. I hope to obtain a better understanding of how research is conducted in real life as well as establish connections that I can use one day whether it be for undergraduate research or for my honors college thesis. I think that research experience applies to my goals because it will teach me to look at things from a scientific standpoint. Having a better understanding of how scientific research works may allow me to conduct my own research as a veterinarian."

**Student #1 Essay #2**-"Overall, I fell that this class greatly enhanced my understanding of the scientific world and how research is conducted. Although it was somewhat more difficult, I feel that this class was a better opportunity than its regular lab counterpart. It was far more interesting to observe and record data that had not been found before as opposed to doing a lab done by thousands of bored students before me. I enjoyed learning lab techniques like the culture transfers as well as how to build presentations for the class. It was very interesting and challenge to go in depth in such a niche area of research, and I feel that it better prepared me for both future lab classes and any research I may do in the future. I struggled with the communication aspect, particularly presenting, but I feel that I improved my public speaking abilities and built more confidence than I had before."

**Student #2 Essay #1**- "I am currently majoring in Human Movement Science with a concentration in Kinesiology Pre-med. As of now, I plan to minor in Biology as I hope to continue my education by attending medical school. After medical school, my goal is to finish as a pediatric anesthesiologist. During my years at [high school], my science classes were always based on the simple science experiments. I never had the chance to learn about real research. When I think about the term "research," the first thing that comes to mind is a lab rat spinning on a wheel and a scientist with big, frizzy hair. As I was receiving emails about my classes before school started, I learned that my … class was a "CURE" lab, and I never heard of "CURE" before. I am very excited to be a part of current research that nobody has ever done before, which also makes me nervous at the same time. I have never done lab research, so jumping straight into a current research project scares me because it may be a lot of hard work for me to handle for my first time. After I attended my first lab, my nerves were gone. I learned that it is okay to mess up because there is no certain outcome that should happen with current research. Once this semester is finished, I hope to expand my excitement about research, and obtain a student job…for my undergraduate research project. I am very confident that the research hours I will have under my belt going into medical school will help me push through and finish with a high amplitude of experience as I earn my Doctorate."

**Student #2 Essay #2**- "I was very intimidated about CURE research. Now that I am going into the last week of this class, I am very honored to have been able to work with it. I have many friends who are enrolled in [the regular section], and all they do is write charts down and work with results that have already been found. Knowing that I could've been stuck with working in those labs makes me even more happy that I was a part of the CURE lab. Working with a new bacterium was a great experience. I definitely believe that this CURE lab has prepared me for all types of research that will come up for my future lab classes. Not only did I learn how to conduct the experiments in the lab, I was able to learn how to create a well-organized and professional poster along with a lab report. I learned the rights and wrongs of presenting in front of people and received feedback of my presentation by being videoed. I would recommend to any biology student to take a CURE lab at least once. It has prepared me for the future and developed my knowledge and skills in ongoing research. After completing every experiment and characterizing LSUCC0135, I truly feel like a researcher, and I have been a part in helping define such a cool bacterium that is very close to my home. I would love to thank CURE for giving me this great opportunity!"

#### Appendix 27: Example student poster

### Characterization of Carbon Substrate Usage, Temperature, and Salinity of Coastal Louisiana Isolate LSUCC0135

Names removed
CURE Students in the Department of Biological Sciences,
Louisiana State University, Baton Rouge, LA, 70808

#### Introduction

- Bacteria are one of the most helpful components of the global ecosystem because of their ability to cycle chemical elements through the atmosphere
- In order to connect the microbial identify to its role in the environment, it is important to test carbon usage, temperature, and salinity through the use of artificial seawater media, MWH3[1]
- By learning which carbon substrates the bacteria generates energy from, we can then locate the same carbon substrates within the environment
- Bacteria does not have the ability to regulate their own internal temperature, so they can only survive within their cardinal temperature
- Cell death will occur when bacteria are not within the appropriate salinity range
- Carbon substrates, temperature, and salinity fluctuate in coastal Louisiana waters due to environmental factors
- LSUCC0135 was isolated from Freshwater City, Louisiana using high throughput dilution-to-extinction culturing[2]

##### Hypothesis

LSUCC0135 will thrive through the use of every carbon substrate tested, in a temperature range of 12˚C to 33˚C, and in a salinity range of 6 to 33.

#### Methods

##### Carbon Substrate Usage

Carbon plate 1 was initially filled with 1.5 mL MWH3, without Carbon. Following sterile technique, each well was inoculated with 5µl of a different substrate and 6.4 µl of LSUCC0135. After sitting for two weeks, cells were counted using Guava flow cytometer. Plate 1, 6.4 µl of LSUCC0135, was transferred to plate 2 after the two weeks; 5 µl of a different carbon substrate was added to each well. The counts and growing time were the same. The method was repeated for the transfer of plate 2 to plate 3.

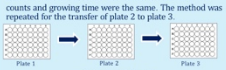

##### Temperature

Sterile technique was followed before beginning. The flask originally contained 50 mL of MWH3, and 200 µl of LSUCC0135 was then added. Flasks were inoculated at either 12˚C, 24˚C, 33˚C, or 40˚C. The flasks were inoculated for a duration of six days while counted daily using Guava flow cytometer.

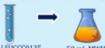

##### Salinity

The salinity experiment began with sterile technique once again. Four medias with different salinities were tested: MWH1 (35) , MWH2 (23), MWH3 (12), and MWH4 (6) were tested. The flasks originally contained 50 mL of the designated media. 200 µl of LSUCC0135. Counts were done daily using Guava cell cytometry.

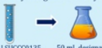

#### Results

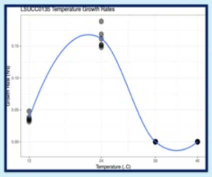

Figure 1. For each temperature tested, growth rates for LSUCC0135 were conducted. The x-axis represents the temperature that were tested in degrees C: 12, 24, 33, and 40. The y-axis represents the growth rates in hours

Table 1. This is a table of the results from the Carbon Substrate method. Under Plate 1-3, the plus sign means that LSUCC0135 grew.

| Carbon Substrate | Plate 1 | Plate 2 | Plate 3 |
|---|---|---|---|
| Arginine | + | + | |
| Cysteine hydrochloride monohydrate | + | + | |
| Histidine | + | + | |
| Isoleucine | + | + | |
| Leucine | + | + | |
| Lysine | + | + | |
| Methionine | + | + | |
| Phenylalanine | + | + | |
| Threonine | + | + | |
| Tryptophan | + | + | |
| Tyrosine | + | + | |
| Valine | + | + | |
| Glutamine | + | + | |
| Dextrose | + | + | |
| Ribose | + | + | |
| Sodium Pyruvate | + | + | |
| Sodium Citrate | + | + | |
| Glutamate | + | + | |
| Sodium Acetate | + | + | |
| Sodium Succinate dibasic hexahydrate | + | + | |
| A-ketoglutaric acid sodium salt | + | + | |
| [illegible] | + | + | |
| Glycine | + | + | |
| Choline Chloride salt | + | + | |
| Cyanate Potassium salt | + | + | |
| Sucrose | + | + | |
| Fructose | + | + | |
| Glucose Solution | + | + | |
| L-Ornithine | + | + | |
| Serine | + | + | |
| Uninoculated b media | + | + | |
| Uninoculated b media | + | + | |
| Uninoculated b media | + | + | |
| Uninoculated b media | + | + | |
| Uninoculated b media | + | + | |
| Uninoculated b media | + | + | |
| Base+LSUCC0135 | + | + | |
| Base+LSUCC0135 | + | + | |
| Base+LSUCC0135 | + | + | |
| Base+LSUCC0135 | + | + | |
| Base+ no LSUCC0135 | + | + | |
| Base+ no LSUCC0135 | + | + | |
| Base+ no LSUCC0135 | + | + | |
| Base+ no LSUCC0135 | + | + | |
| MWH3+LSUCC0135 | + | + | |
| MWH3+LSUCC0135 | + | + | |
| MWH3+LSUCC0135 | + | + | |
| MWH3+LSUCC0135 | + | + | |

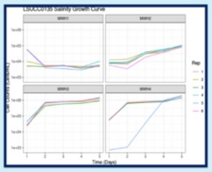

Figure 3. Growth curves of LSUCC0135 for each salinity level tested, which had a range from 6-35. MWH1 was 35, MWH2 was 23, MWH3 was 12, and MWH4 was 6. The x-axis represents time in days for 6 days, and the y-axis represent the Cell Counts in cells/mL. Each colored line represents a replicate.

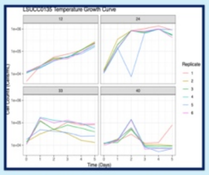

Figure 2. Results for the experiments of the four temperatures tested. Each color represents a replicate that was put into four different temperature flasks. On the x-axis are the days that was tested with the y-axis telling us the cell count for each day.

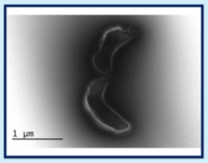

Figure 4. This is a TEM picture of LSUCC0135

According to Table 1, LSUCC0135 used every carbon substrate as an energy source. The growth rate for LSUCC0135 increased as the temperature increased from 12˚ C to 24˚ C (Figure 1). According to Figure 2, the minimum temperature for LSUCC0135 is unknown because it is out of the temperature range that we tested. Cell death will occur below 12˚ C, therefore; further testing needs to be done to find an exact minimum. The maximum temperature is 40˚ C with an optimum of 24˚ C. In the flask with media at 24˚ C, the cells reached maximum growth producing the highest cell count. According to figure 3, the cell counts were low for the salinity level 35, MWH1, which means that the salt concentration was too high. For MWH2, the cell counts steadily increased, and continued to increase for MWH3, which has a salinity level of 12. The cell counts were extremely similar for MWH4 except for replicate 5. From day 1 to day 2, the cell counts were low, and shot up to equal the cell counts for the other replicates for day 4 to day 6. The optimum salinity level for LSUCC0135 is 12, which is MWH3.

#### Discussion

- Results of the experiment partially support the hypothesis by LSUCC0135 using all carbon substrates as an energy source, surviving in a range of temperatures from ~9˚C to ~30˚C, and thriving in a salinity range of about 6 to 30
- There is a huge possibility LSUCC0135 is an autotroph because it is highly unlikely it is able to consume all carbon substrates tested and the control with no carbon was positive
- LSUCC0135 has the capability to be prevalent through a moderate climate and a key component of the ecosystem, but could not survive in an extreme environment. Louisiana is not an extreme environment, so this makes sense.
- The bacterium's role in the carbon cycle is questionable since LSUCC0135 does not depend on organic carbon as an energy source.
- LSUCC0135 could have a key role in the maintenance of the coastal Louisiana ecosystem because it does not depend on the availability of the organic carbon

#### Future Directions

- In the next set of experiments, the exact minimum and maximum temperatures should be found, as well as the exact minimum and maximum salinity for LSUCC0135
- The pressure range for the organism should also be found to narrow down the possible depths LSUCC0135 could be prevalent in
- The amount of nitrogen that bacteria convert within the nitrogen cycle determines the amount that plants are able consume from the soil, so LSUCC0135's nitrogen intake should be found

#### Acknowledgements

We would like to thank LSU CURE, Dr. Thrash, Celeste Lanclos, Mindy Brooks, and Dr. Christopher Gregg for the instruction, support, and funding throughout the duration of our experiment.
